# Cryo-EM led analysis of open and closed conformations of Chagas vaccine candidate TcPOP and its antibody response characterisation

**DOI:** 10.1101/2024.03.26.586384

**Authors:** Sagar Batra, Francisco Olmo, Timothy J Ragan, Merve Kaplan, Valeria Calvaresi, Asger Meldgaard Frank, Claudia Lancey, Mahya Assadipapari, Cuifeng Ying, Weston B. Struwe, Emma Hesketh, John M. Kelly, Lea Barfod, Ivan Campeotto

**Affiliations:** School of Biosciences, Division of Microbiology, Brewing and Biotechnology, University of Nottingham, Sutton Bonington Campus, Loughborough, LE12 5RD, UK; Interdisciplinary Biomedical research Center, School of Science and Technology, Nottingham Trent University, NG11 8NS, UK; Department of Parasitology, Faculty of Sciences, University of Granada, Granada, Spain; Leicester Institute of Structural and Chemical Biology, University of Leicester, Lancaster Road, Leicester, LE1 7RH, UK; Physical and Theoretical Chemistry Laboratory, Department of Chemistry, University of Oxford, South Parks Road, OX1 3TA, UK; The Kavli Institute for Nanoscience Discovery, Dorothy Crowfoot Hodgkin Building,University of Oxford, OX1 3QU, UK; Department of Immunology and Microbiology, Centre for Medical Parasitology, Faculty of Health and Medical Sciences, University of Copenhagen, Copenhagen, DK; Advanced Optics & Photonics Laboratory, Department of Engineering, School of Science & Technology, Nottingham Trent University, Nottingham NG11 8NS, UK; Department of Infection Biology, Faculty of Infectious and Tropical Diseases, London School of Hygiene and Tropical Medicine, Keppel Street, London, WC1E 7HT

**Author notes:** equal author contribution (Sagar Batra, Francisco Olmo).

**Keywords:** Chagas disease, cryo-EM, enzymology, SAX, MD, plasmonic optical tweezers, HDX-MS, neutralizing antibodies

## Abstract

Chagas disease, caused by the protozoan parasite *Trypanosoma cruzi*, remains a significant global public health concern. Despite its profound health impact in both endemic and non-endemic areas, no vaccine is available, and the existing therapies are outdated, producing severe side effects. The 80kDa prolyl oligopeptidase of *Trypanosoma cruzi* (TcPOP) has been recently identified as a leading candidate for Chagas vaccine development. We report the first three-dimensional structure of TcPOP in open and closed conformation, at a global resolution of 3.8 and 3.6 Å respectively, determined using single-particle cryo-electron microscopy. Multiple conformations were observed and further characterized using plasmonic optical tweezers and hydrogen-deuterium exchange mass spectrometry. To assess the immunogenic potential of TcPOP, we immunized mice and evaluated both polyclonal and monoclonal responses against the TcPOP antigen and its homologues. The results revealed invasion blocking properties of anti-TcPOP polyclonal response via parasite lysis and its cross-reactivity versus closely-related POPs but not with human homologues. We were also able to produce and characterise three monoclonal antibodies, one of which showed neutralising properties and inhibition of parasite invasion via non-lytic mechanism. Collectively, our findings provide critical structural and functional insights necessary to understand the immunogenicity of TcPOP for future Chagas vaccine development and diagnostic applications.

## INTRODUCTION

Chagas disease is a chronic life-threatening parasitic disease caused by *Trypanosoma cruzi* and represents a significant health burden in 21 countries of Latin America^1^. Chagas continues to expand beyond endemic zones because of human migration and global warming^2^, with 6 million infected people worldwide, leading to approximately 12,000 deaths per year (PANHO ^3^). The major transmission route is via the bite of insects belonging to the Triatomine species bug in endemic regions, although other routes include congenital transmission, organ transplants, blood transfusion or oral transmission ^4^.

Chagas disease becomes symptomatic during the chronic stage when muscle cells in the heart and gut are compromised by both infection and immune responses against the parasite; this results in cumulative damage that leads to severe cardiac complications and heart failure in approximately 30% of patients, with an estimated global economic hardship of $7.19 billion US per year^5^. Chagas is therefore, one of the major global neglected tropical diseases (NTDs). There is no vaccine, and current therapy relies on the usage of the drugs nifurtimox and benznidazole^6^, which can cause severe side effects including sterility, blindness, and deleterious effects in adrenal, colon, oesophageal and mammary tissue ^7^. Additionally, therapies are more effective during the acute phase of the disease, and less so during chronic phase. Infection with *T. cruzi* is generally considered to be life-long, therefore representing a “time bomb” on the health systems around the world.

To complicate the epidemiological scenario, due to the evolutionary plasticity of *T.cruzi*, there are six discrete typing units (DTUs)^8^ found to infect humans. An additional DTU has been identified in bats (Tcbat), although transmission to humans has not been yet reported^9^. DTUs differ in geographical and ethnic distribution, clinical manifestations, and reservoir hosts, which include more than 150 species of mammals ^10^.

Diagnostic tools are often strain specific and unable to detect congenital transmission in newborn babies for the first 6 months, due to passive immunity from the mother, which can cause life-threatening complications later in life (PANHO^11^). Amidst these challenges, a member of the endopeptidase enzyme family, *T. cruzi* 80kDa prolyl-oligopeptidase (TcPOP, enzyme ID EC 3.4.21.26), has been identified as novel vaccine target for Chagas ^12^. TcPOP is expressed in both the extracellular blood trypomastigote and the replicative intracellular amastigote ^13^. It degrades collagen and fibronectin extra-cellular matrix components on host cells, facilitating parasite invasion. Its secretion into the blood during invasion, expression throughout the mammalian stages of the life cycle, and the more than 98–99% sequence conservation across DTUs ^12^ makes it an ideal candidate for vaccine development. This has been further supported in a murine model, as polyclonal antibodies against recombinant TcPOP, produced in *E. coli*, protected the mice from a lethal dose of the parasite^12^. *T. cruzi* triggers a complex immune response in humans, involving both innate and adaptive systems. Initially, the innate response is activated, including cytokine production by phagocytes and natural killer cells, which produce interferon-gamma ^14^. The adaptive response involves T and B lymphocytes, with T cells orchestrating cellular immunity and B cells producing antibodies, although these are often ineffective due to the parasite’s evasion strategies ^15^. *T. cruzi* employs mechanisms including hijacking of the TGF-β signalling pathway and modulating immune responses to evade detection and establish a chronic infection. Achieving a potent and protective humoral response would be an ideal approach to block transmission in humans ^16^.

No structure of parasite prolyl oligopeptidases (POPs) exists to date, despite POPs being reported in *Leishmania infantum (*LiPOP,^17^), *Trypanosoma brucei (*Tb*POP)*^18^ and *Schistosoma mansoni (*SmPOP)^19^. The POP family is widely distributed across Eukaryotes and Prokayotes ^20^. The closest protein homologues to TcPOP for which experimental structures are available are from *Haliotis discus hannai* (PDB code 6JCI), porcine muscle POP (PDB code 1QFM), *Pyrococcus furiosus* (PDB code 5T88) and Human POP (PDB 3DDU). Comparative homology modelling studies of TcPOP have been based on porcine POP^21^ and predict a cylindrical-shaped structure, consisting of a peptidase domain and a seven-bladed β-propeller domain with the substrate binding site and the canonical Asp, Ser, His catalytic triad located in the middle of the two domains ^22^.

The TcPOP sequence is highly similar to LiPOP (63%), TbPOP (73%) and HuPOP (43%). We therefore also expressed these other POPs and immunised mice against TcPOP to investigate cross-reactivity related to epitope conservation across species. Polyclonal antibodies (pAbs) were isolated and characterised from mice and showed striking parasite neutralising properties in cell-invasion assays. Additionally, one monoclonal antibody was also identified from the mice exhibiting a neutralising response. The structure of TcPOP was solved by cryo-EM in closed and open conformations, revealing unexpected details on its dynamicity, corroborated by a plethora of in solution biophysical and structural techniques, which offers insights into the dynamicity of TcPOP,paving the way for new therapeutic interventions against Chagas disease.

## MATERIALS AND METHODS

### Bioinformatic analysis of TcPOP and TcPOP homologues

Protein sequences were extracted from the Uniprot database ^23^ for TcPOP (Q71MD6), TbPOP (Q38AG2), LiPOP (A4ICB5) and HuPOP (P48147) and aligned in BLASTP ^24^ to retrieve the top 100 homologues of TcPOP and a phylogenetic tree was built in iTOL ^25^ after MSA generation in MUSCLE ^26^. Comparative homology models were obtained with SWISS-MODEL ^27^ and when it became available to the scientific community, AlphaFold2 was used to obtain AI-based models ^28^. Phylogenetic analysis and sequence conservation mapping on TcPOP was done using CONSURF ^29^. The topology diagram of TcPOP was created using PDBsum ^30^. Geometry of the models was assessed with MOLPROBITY ^31^ and 3D alignment performed in PyMOL ^32^.

### Expression and purification of POPs by bacterial fermentation

The codon-optimised genes encoding for TcPOP and orthologues were purchased from TWIST and inserted in the pET28(a)+ vector (Novagen). The resulting N-ter and C-ter His_6_-tagged proteins were all recombinantly expressed and purified from the *E. coli* NiCo21(DE3) strain (NEB). Cells were grown at 37°C supplemented with 2XYT medium (Melford) until OD_600nm_ at 0.6-0.8 was reached. Protein expression was induced using 0.5 mM 1-thio-β-D-galactopyranoside (IPTG, Generon) using an 8L *in-situ* bioreactor (INFORS – Techfors S) at 100 rpm, for 18 hrs at 20°C with pH control at 7.0 (+/− 0.1), resulting in 16 grams of bacterial cell pellet. Lysis was performed using BugBuster (Nalgene) and lysate was clarified by spinning at 50,000xg for 30 min at 4°C. Affinity chromatography was performed using Co^+2^-NTA resin (Thermo Fisher). Washing buffer (50 mM sodium phosphate, 500 mM NaCl, 30 mM imidazole, pH 7.4) and elution buffer (50 mM sodium phosphate, 500 mM NaCl, 500 mM imidazole, pH 7.4) were used during affinity purification, followed by dialysis with 3.5 kDa MWCO Dialysis membranes (Thermo Fisher) at 4°C for 18 hrs against 20 mM Hepes and 150 mM NaCl, pH 7.4. Finally, the proteins were concentrated using 30 kDa MWCO Amicon (Millipore) ultra-centrifugal filters to 10 mg/mL and immediately injected into gel filtration column S200 10/300 (Cytiva) equilibrated with 20 mM Hepes and 150 mM NaCl, pH 7.4. Elution fractions containing protein were pooled for further characterisation.

### Western blot analysis

Samples were assessed on the SDS-PAGE gel (Bolt™ Bis-Tris Plus Mini Protein Gels, 4-12% gradient) (Page Ruler Pre-stained Ladder, ThermoFisher) and transferred to nitrocellulose membrane using the Trans-Blot Turbo system (Biorad). Nitrocellulose membrane was incubated with His_6_-tagged Monoclonal Antibody - HRP (Invitrogen) with 1:3000 v/v dilution in PBS-T, followed by the signal detection using SuperSignal West Pico PLUS Chemiluminescent Substrate (Cat. 34580, ThermoFisher) in an iBright system (Thermo Fisher).

### Differential scanning fluorimetry

Stability measurement of prolyl oligopeptidases was carried out using a fluorescence assisted thermal unfolding assay. A fluorescence stain SYPRO Orange dye solution (ThermoFisher) was used and diluted to 5x concentration in 20 mM Hepes and 150 mM NaCl, pH 7.4. The assay was performed with a final protein concentration of 1 µM in a total volume of 20 µL. The temperature of the protein samples was gradually increased from 25°C to 95°C at a rate of 5°C per min, using the Rotor-Gene Q (Qiagen). Lastly, the data were analysed using non-linear regression method to determine the midpoint temperatures (T_m_) of the thermal shift.

### Enzymatic tests for POPs

Fluorogenic POP substrate Suc-Gly-Pro-Leu-Gly-Pro-7-amido-4-methylcoumarin (AMC) (HANGZHOU JHECHEM CO LTD) was diluted in PBS to 1-25 µM final concentrations in 96-well plates (ThermoFisher) to measure enzyme activity parameters in real time, with end-point reactions at 10 and 30 mins, using a Gen5™ Microplate Reader and Imager Software. Stock solutions were made in PBS and stored in aliquots at –20°C. The microplate readings were performed at excitation 360/40 nm and emission 460/40nm using BioTek Synergy LX Multimode plate reader.

### Mice immunisations

The conducted animal research strictly conformed to the standards delineated by the Federation of European Laboratory Animal Science Associations (FELASA). Ethical clearance for the experimental methodologies was granted by the Danish Animal Experiment Inspectorate, as indicated by their approval number 2018-15-0201-01541. For the purpose of immunization, female BALB/c ByJR mice, aged six weeks, were obtained from Janvier Labs. Three mice were immunized intramuscularly with 20 µg TcPOP, emulsified in 50% v/v AddaVax adjuvant (InvivoGen). This was followed by two additional intramuscular injections at biweekly intervals. A concluding intraperitoneal injection of 20 µg TcPOP in PBS was carried out two weeks post the last boost. Three days after the final injection, the mice were humanely euthanized for the extraction of spleen and blood from which sera were subsequently obtained.

### Production, purification and conjugation of anti-TcPOP mAbs

After picking the best immune responders, hybridoma cell lines were generated by fusing splenocytes of immunized mice with myeloma cells (ClonaCell-HY hybridoma cloning kit)^33^. Hybridoma cell lines were harvested 14 days after the fusion and plated into 96-well culture plates in HT supplemented media. Screening of hybridoma cell lines producing antibodies specific to TcPOP was performed by ELISA, as described below. Monoclonal TcPOP-specific hybridoma cell lines were obtained by single cell sorting using the FACSMelody (BD). For large-scale mAb production, hybridoma cell lines were cultured in 4×250 mL cell culture flasks (Corning) as per manufacturer’s instructions. Monoclonal antibodies were purified by affinity chromatography using a 5 mL protein G sepharose column (Cytiva) on an ÄKTAxpress system (Cytiva). Antibodies were eluted at 0.8 mg/mL in 0.1 M glycine buffer, pH 2.8 and immediately neutralized with 1/10 v/v 1 M Trizma hydrochloride solution (Sigma Aldrich), pH 9.0 to obtain the final pH 7.4. Buffer exchange to 1 x PBS was performed using a desalting column (Generon) and eluted protein was concentrated with 30 kDa MWCO Amicon (Millipore) to 20 mg/mL. The 3 mAbs were specifically conjugated to HRP using an EZ-*Li*nk™ Plus Activated Peroxidase Kit (ThermoFisher) and detected with ECL (Pierce) using the iBright system.

### Enzyme-*Li*nked Immunosorbent Assay (ELISA)

Antibody-producing hybridoma cells against TcPOP were ascertained through Enzyme-*Li*nked Immunosorbent Assay (ELISA). In summary, MaxiSorp flat-bottom 96-well ELISA plates (ThermoFisher) were coated with recombinant TcPOP (2 µg/mL in PBS) overnight at 4°C under shaking. Plates were there washed with PBS supplemented with 0.05% v/v Tween20 (PBS-T) and blocking performed for 1 hr with casein blocking solution (Pierce). The blocking solution was then removed and replaced with 50 µL of hybridoma supernatant and incubated for 1 hr at RT under shaking. Plates were washed three times with PBS-T before undergoing incubation with 1:10000 v/v anti-mouse IgG (γ-chain specific) for 1 hr, followed by three 5 min washes with PBS-T. The positive wells were identified by adding TMB plus2 (Kementec) for 20 mins and quenched using 0.2N sulfuric acid. Colorimetric and absorbance signals were measured at 450 nm. Data were analysed using GraphPad (Prism).

### Biological studies

#### Cell and parasite culture

COLO-N680 cells (human oesophageal squamous cell carcinoma line) were maintained in complete MEM medium, consisting of Minimal Essential Medium (Sigma) supplemented with 5% (v/v) heat-inactivated foetal bovine serum (hiFBS, Cytiva), 100 U/ml penicillin, and 100 μg/ml streptomycin. Cells were incubated at 37°C in a 5% CO_2_ atmosphere and sub-cultured every 3 days at a 1:5 ratio. COLO-N680 cells were seeded in complete MEM at 80-90% confluency in T25 vented flasks and infected with 2 × 10^6^ Tissue Culture Trypomastigotes (TCTs) derived from previously infected cells. Infected cultures were maintained for 5 days post-infection. Free-swimming TCTs were isolated by collecting and centrifuging the culture medium at 1600xg. The resulting pellets were resuspended in Dulbecco’s Modified Eagle Medium (DMEM) containing 5% hiFBS and maintained at 37°C for up to 4 hrs before use. Motile trypomastigotes were quantified using a hemacytometer.

#### *In vitro* neutralization cell-based invasion assay

Trypomastigotes obtained as above were incubated in plain DMEM for at least 30 mins before starting the assays. Then parasites were pelleted and incubated in DMEM prepared with the sera or monoclonal antibody suspension, according to the concentration assayed. They were incubated for 4 hrs at 37°C in an orbital shaker, unless specified differently. Then parasites were washed twice in DMEM to remove the sera/antibodies from the medium and resuspended in DMEM prior to infection. Cells were infected at a multiplicity of infection of 10:1 (parasite:cell). After 4 hrs of infection, cells were washed three times with PBS to remove non-internalized parasites and incubated with fresh complete MEM for 72 hrs to allow intracellular amastigote replication. Then cells were stained with 5 µg/ml of Hoechst and live images were acquired using an inverted Nikon Eclipse T2i epifluorescence microscope. Infected cells were detached using 1 mL TrypLE Express (Gibco™) for 12 mins at 37°C and fixed with 4% paraformaldehyde (PFA) for 1 hr. Cells were then washed in PBS by centrifugation and resuspended in flow cytometry staining buffer (FCSB) and flowed in an Attune NxT Flow Cytometer (Thermo Fisher). Gating was performed using a non-infected culture as a control.

#### Immunofluorescence assays

For the live binding assays, parasites were immediately fixed with 4% (v/v) formaldehyde for 1 hour at room temperature following the indicated incubation time with antisera or monoclonal antibody suspensions. Fixed parasites were washed twice with phosphate-buffered saline (PBS) and allowed to settle by gravity onto 10-well slides for immunofluorescence (MP Biomedicals™). Parasites were then permeabilized and blocked in 0.5% saponin containing 10% donkey serum in PBS for a minimum of 1 hour, followed by three washes with PBS, each for 5 minutes. For classic immunostaining, where parasites were directly fixed after isolation from culture, an additional incubation step was performed: parasites were incubated at room temperature for 3 hours with either a 1:50 dilution of polyclonal mouse antiserum or a suspension of monoclonal antibody at 1000 µg/ml in 0.5% saponin containing 1% donkey serum in PBS. After three washes with PBS, each for 5 minutes with agitation, parasites were incubated with a secondary goat anti-mouse antibody (Alexa Fluor 488, Invitrogen) for 1 hour at room temperature, protected from light. Slides underwent a final series of three washes with PBS, each for 5 minutes, and were then allowed to air-dry. Finally, slides were mounted using Vectashield containing DAPI for DNA staining. Images were acquired using an inverted Nikon Eclipse T2i epifluorescence microscope and processed using the NIS-Nikon software.

### Biophysical studies

#### Binding kinetics of TcPOP and anti-TcPOP mAbs using biolayer interferometry

Binding of anti-TcPOP mAbs to TcPOP were measured by kinetic experiments carried out on an Octet R4 (Sartorius). All samples were buffer exchanged into Sartorius Kinetics Buffer, according to the manufacturer’s instructions. All measurements were performed at 200 μL per well in Sartorius kinetic buffer at 25°C in 96-well black plates (Greiner Bio-One, Cat# 655209). ProG (Cat. Nos. 18-5082, 18-5083, 18-5084) were used to immobilize anti-TcPOP mAbs for 1800s. Immunogens were four-fold serially diluted in kinetic buffer in the range of 64 nM to 4 nM. Assays were performed in three sequential steps with Octet BLI Discovery 12.2.2.20 software (Sartorius): Step 1, biosensor hydration and equilibration (300s); Step 2, immobilization of anti-TcPOP IgG1 mAbs on a ProG biosensor (600s); Step 3, wash and establish baseline (60s); Step 4, measure TcPOP association kinetics (1800s); and Step 5, measure TcPOP dissociation kinetics (600s). The acquired raw data for the binding of anti-TcPOP mAbs with TcPOP were processed and globally fitted to a 1:1 binding model. Binding kinetics measurements were conducted in triplicate, and reported values represent the average. Data were analysed using Octet Analysis Studio 12.2.2.26 Software (Sartorius) and graphs produced using GraphPad (Prism).

#### Epitope binning studies using Bio-layer interferometry (BLI)

Anti-TcPOP mAb2 and anti-TcPOP mAb3 were applied sequentially to assess competition using the BLI. Assays were performed in seven sequential steps with Octet® BLI Discovery 12.2.2.20 software (Sartorius): Step 1, biosensor hydration and equilibration (300s); Step 2, immobilization of TcPOP NiNTA biosensors (600s); Step 3, wash and establish baseline (60s); Step 4, measure anti-TcPOP mAb2 association kinetics (1800s); Step 5, measure anti-TcPOP mAb2 dissociation kinetics (600s); Step 6, measure anti-TcPOP mAb3 association kinetics (1800s); and Step 7, measure anti-TcPOP mAb3 dissociation kinetics (600s). The acquired raw data for the binding of anti-TcPOP mAbs with TcPOP were processed and globally fitted to a 1:1 binding model with Octet Analysis Studio 12.2.2.26 Software (Sartorius). The binding kinetics measurements were carried out in three replicates. Values reported are the average among triplicates.

#### Estimation of apo-TcPOP using SAXS

SAXS was performed at the B21 beamline (Diamond *Li*ght Source, Oxon, UK). TcPOP was buffer exchanged into 20 mM HEPES and 150 mM NaCl (pH 7.4) at 277K before data collection. Using an Agilent 1200 HPLC system, 50 μL of TcPOP at 8 mg/mL was loaded onto a superdex S200 3.2/300 column for SEC-SAXS. Also, static-SAXS was performed separately at different concentrations extrapolated to zero (from 6 mg/ml to 0 mg/ml, i.e buffer condition). For SEC-SAXS, X-ray intensity data were collected as the eluent moved from the column to the beam at a flow rate of 0.16 mL/min to collect 600 frames at 3 sec intervals, while static SAXS samples were exposed to X-rays to collect 21 frames at 1 sec time intervals. The intensity was plotted against its angular dependants q (q = 4πsinθ/λ) while, the system operated with an exposure time of 3 sec at 12.4 keV (1Å) using a EIGER 4 M detector. Data were analyzed using the BioXTAS RAW ^34^, ATSAS program suites ^35^, DENNS^36^ and plotted using GNOM^37^.

### Mass photometry

#### Buffer optimisation for TcPOP

Mass photometry (MP) experiments were conducted using the Refeyn OneMP mass photometer, after cleaning coverslips and gaskets with 100% isopropanol and water. Measurements, performed in triplicate, involved systematic optimizations of pH values in the range 7.0-8.0 and with NaCl concentration range 50-300mM within a 20 mM BTP buffer. Protein was diluted in buffer to a final concentration of 120 nM into a gasket well, followed by focal point acquisition and data analysis using Refeyn AcquireMP 2.3.1 software. MP movies (6000 frames, 20 frames per sec) were captured within a 10.8 × 10.8 μm field and processed with Refeyn DiscoverMP 2.3.0 software. Robust data analysis ensued, leveraging a contrast-to-mass (C2M) calibration approach. Calibration involved introducing 3 μL of a 1:100 v/v pre-diluted NativeMark standard (LC0725, Thermo Scientific) to an acquisition well, yielding masses (66, 146, 480, 1048 kDa) that informed the calibration curve employed in DiscoverMP software. Experiments were performed at the Leicester Institute for Structural and Chemical Biology (LISCB, University of Leicester, UK).

#### Anti-TcPOP binding measurements

MP measurements were conducted using a Refeyn TwoMP (Refeyn Ltd) as previously described ^38^. Briefly, glass coverslips (High Precision No. 1.5H, Marienfeld Superior) were cleaned by sequential sonication with Milli-Q H_2_O, 50% isopropanol and again Milli-Q H_2_O. Cleaned coverslips were dried using nitrogen flow. CultureWell^TM^ reusable gaskets (3 mm diameter x 1 mm depth, Grace Bio-Labs) were used to assemble sample chambers. Coverslips were placed on the MP sample stage and a single gasket was filled with 20 μL DPBS (wo/ calcium and magnesium, pH 7.4, ThermoFisher Scientific) to find focus. TcPOP, mAb1, mAb2 and mAb3 were measured separately at a final concentration of 20 nM. For TcPOP-antibody binding assays, 5 μM TcPOP was mixed with 5 μM mAb2 or mAb3 in a 1.5 mL Eppendorf tube at a 1:1 v/v ratio. The sample mixture was equilibrated for 10 mins and diluted 1:100 before data acquisition. Acquisition settings were adjusted within AcquireMP (2023 R1.1, Refeyn Ltd) as a large field of view, frame binning = 2, frame rate = 128.2 Hz, pixel binning = 6, exposure time = 7.65 ms. Movies were taken over 60 sec. Mass calibration was performed using an in-house protein standard including 90-720 kDa oligomers. Data were analysed and histograms were created with Discover MP (v2023 R1.2, Refeyn Ltd). Experiments were performed at the New Biochemistry building (University of Oxford, UK).

#### MD simulations

Atomics coordinates of TcPOP were retrieved from the AlphaFold database. To calculate the conformational dynamics of TcPOP, all-atom molecular dynamics simulations were conducted on an Ada High Performance Computer (HPC, University of Nottingham) using the GROMACS 2021.2-fosscuda-2020b package. GROMOS 54a7 forefield was applied and hydrogen atoms were incorporated using the pdb2gmx module, and topology files were generated under periodic boundary conditions (PBC) employing a cubic periodic cell. The protein was centrally placed, solvated using simple point charge (SPC) 216 water molecules and positioned 1 nm from the edges, with NaCl counter ions added for system neutralization.

Following energy minimization, the canonical ensemble (NVT) underwent equilibration for 100 psec without pressure coupling and Berendsen thermostat was initially applied. Subsequently, temperature (298 K) was maintained by velocity rescaling with a stochastic term, while the isothermal-isobaric ensemble (NPT) with a 1 bar pressure for 100 psec, using the Parrinello–Rahman method, was implemented. The LINCS algorithm constrained H-bonds, and the MD simulations ran for 500 nsec with a 2-fs time step. The resulting trajectory was analysed using inbuilt functions of the GROMACS package^39^.

### Cryo-EM sample preparation, data collection and processing

#### Cryo-EM grid preparation

Homogeneous samples from SEC purified in 20 mM HEPES, 150 mM NaCl, pH 7.4 were freshly used to prepare the grids. Fraction corresponding to the SEC peak at 15 mL (Fig.S2) was used at a final concentration of 0.2 mg/mL. Firstly, cryoEM grids, R1.2/1.3 carbon, Au 300 (Quantifoil), were glow discharged in the presence of amylamine for 30 sec at 10 mA on a Quorum GloQube glow-discharge unit. Four microliters of the freshly prepared TcPOP sample were applied to the grid and blotted for 3 sec, with blot force 10, prior to flash-cooling in liquid ethane using a Vitrobot Mark IV (FEI ThermoFisher), set at 4 °C and 100% humidity.

#### CryoEM data collection

CryoEM grids were imaged using a 300 KeV Titan Krios G3 (ThermoFisher Scientific) transmission electron microscope (Midlands Regional Cryo-EM Facility, University of Leicester) at a calibrated pixel size of 0.656 Å. Electron micrographs were recorded using a K3/GIF (Gatan Imaging Filter) direct electron detector (Gatan Inc.) and EPU automated data acquisition software (ThermoFisher Scientific). Micrograph movies were recorded with 75 fractions, in super resolution, binned by 2 and a total dose of 77 e^-^/pix (dose rate of 15 e-/pix/s), To improve the distribution of particle views, data were collected at 0°, 30° and 35°. At 0° tilt (Dataset 1), the defocus range was collected between −2.3 and −0.8 µm, in regular intervals. At 30° and 35° tilt (Dataset 2), the defocus was set to −1.2 µm.

#### Cryo-EM image processing

Image processing was carried out on the cryo-EM computational cluster at the University of Leicester. Micrographs were pre-processed using Relion5-beta,^40^ sample motion during acquisition was corrected using RELION’s own implementation, Dataset 2 was dose-weighted, Dataset 1 was not. Micrographs were then CTF estimated using CTFFIND4.1^41^. Initially each dataset (i.e. 0° tilt, 30° tilt and 35°) was processed separately. In brief, particles were picked using Topaz^42^, then extracted with a box size – 256pixels (corresponding to 168Å) binned by 4 to 2.624 Å/pix. Particles were filtered in 2D to produce a set of images which were then used for *ab-initio* model generation in RELION 5-beta. Analysis of the 3D initial models revealed “open” and “closed” conformations. The full set of Topaz picked particles was classified in 3D with Blush regularisation^43^ and the classes were inspected and subcategorised into open and closed particles. Re-extraction of the particles at 1.312 Å/pix followed by 3D auto-refinement and CTF parameter refinement of all three datasets was then carried out. Despite having a 3DFSC sphericity > 0.9, The 0° tilt dataset showed severe preferential orientation and concomitant overestimation of resolution and B-factor parameters was observed. All three datasets were then merged but this resulted in only marginally improved maps.

Further analysis of the 35° tilt dataset was then performed with DynaMight^44^. This showed that both the open and closed conformations had a moderate degree of continuous motion, explaining why the map quality was limited despite having a large number of high-quality particles. The consensus deformed back projection of both open and closed conformations were then calculated. Map sharpening was done with Relion’s automatic B-factor estimation as well as DeepEMhancer^45^.

**Table 1.** Cryo-EM data collection, refinement and validation statistics.

| Data collection and processing | Dataset 1 (untilted) | Dataset 2 (tilted) |
| --- | --- | --- |
| Magnification (nominal) | 130,000 x | 130,000 x |
| Voltage (kV) | 300 | 300 |
| Stage (Z) tilt | 0 | -30 and -35 |
| Aberration free image shift (AFIS) | Yes (12 $\mu\text{m}$) | No |
| Electron exposure ($\text{e}^-/\text{\AA}^2$) | 76.43 | 77.59 |
| Illuminated area ($\mu\text{m}$) | 0.65 | 0.57 |
| Fractionation | 75 | 75 |
| Defocus range ($\mu\text{m}$) | -2.3 to -0.8 | -1.2$\mu\text{m}$ |
| Pixel size ($\text{\AA}$) | 0.656 | 0.656 |
| Total micrographs (no.) | 20,080 | 2,292 |
| Total particle images (no.) | 45,400,101 | 2,525,462 |
| Final particle images (no.) | 847,560 (closed)<br>518,852 (open) | 163,153 (closed)<br>126,459 (open) |
| AutoRefine map resolution @ 0.143 | 3.29$\text{\AA}$ (closed)<br>3.50$\text{\AA}$ (open) | 4.00$\text{\AA}$ (closed)<br>4.10$\text{\AA}$ (open) |
| AutoRefine 3DFSC global resolution @ 0.143 (sphericity) | 3.29$\text{\AA}$ (0.929) (closed)<br>3.43$\text{\AA}$ (0.908) (open) | 3.82$\text{\AA}$ (0.929) (closed)<br>4.10$\text{\AA}$ (0.912) (open) |
| DynaMight map resolution @ 0.143 | n/a | 3.57$\text{\AA}$ (closed)<br>3.82$\text{\AA}$ (open) |
| DynaMight 3DFSC global resolution @ 0.143 (sphericity) | n/a | 3.52$\text{\AA}$ (0.960) (closed)<br>3.93$\text{\AA}$ (0.964) (open) |
| Initial model used | *Ab initio* | *Ab initio* |

### Cryo-EM model fitting and refinement

The coordinates of the open and closed conformations of TcPOP were docked in the respective cryo-EM maps using phenix.*dock_in_ma*p^46^. Both maps were refined in PHENIX using real space refinement with 3 cycles of simulated annealing (SA), followed by re-build in place and 10 cycles of refinement at each SA step. Model building was performed in COOT^47^. Structures were validated using the Molprobity tool, as implemented in PHENIX. ChimeraX^48^ was used to generate visual molecular graphics. The FSC of the closed and open CryoEM maps were calculated using EBML-EBI FSC-server^49^. Cryo-EM reconstructions have been deposited in the EM Data Bank (EMDB).

#### Plasmonic Optical nanotweezers: samples preparation and data collection

We used a plasmonic optical tweezers setup which is a modified modular optical tweezers system (OTKB/M, Thorlabs) in the Advanced Optics and Photonics Lab at Nottingham Trent University (Nottingham, UK) with a 852 nm Fabry-Perot laser diode (FPL852S, Thorlabs)^50^. The laser beam was polarised perpendicular to the centre-to-centre line of two circles of the double nanohole (DNH) structure by using a polariser and a half-wave plate and was collimated and focused on the DNH by a 60X air objective (NA 0.85, Nikon). All trappings were performed at a laser power of 25 mW^51^ The transmitted laser intensity was then converted to a voltage signal via a silicon avalanche photodiode (APD120A, Thorlabs) and recorded by a data acquisition card with a sampling rate of 1 MHz. The recorded voltage data (transmission traces) were normalised and filtered using in-house MATLAB scripts, which included a zero-phase Gaussian low-pass filter (MATLAB *filtfilt.m*) with cut-off frequencies of 10 kHz, 1 kHz, 100 Hz, and 10 Hz. Probability density function (PDF) was calculated by using MATLAB function *ksdensity.m*. More information on this specific set-up have been previously reported^52^. SEC Purified TcPOP was used at 1 µM concentration in PBS. DNHs were sealed into the flow cell using cover glass with a double-sided tape as a spacer, providing a microfluidic channel with 3.5 μL volume. The solution was delivered to the flow cell using a 12-way valve and a syringe pump. Initially, TcPOP was infused in the chamber to achieve trapping followed by the sequential infusions of 1 µM, 10 µM and 100 μM substrate Suc-Gly-Pro-Leu-Gly-Pro-AMC to the flow cell at a flow rate of 2 µL/min.

#### Hydrogen-deuterium exchange (HDX) mass spectrometry (MS)

Prior to conducting HDX-MS experiments, peptides were identified by digesting TcPOP using the same protocol and identical liquid chromatographic (LC) gradient as detailed below and performing MS^E^ analysis with a Synapt G2-Si mass spectrometer (Waters), applying collision energy ramping from 20 to 30 kV. Sodium iodide was used for calibration and leucine enkephalin was applied for mass accuracy correction. MS^E^ runs were analysed with ProteinLynx Global Server (PLGS) 3.0 (Waters) and peptides identified in 3 out of 4 runs, with at least 0.2 fragments per amino acid (at least 2 fragments in total) and at least 1 consecutive product, with mass error below 7 ppm were selected in DynamX 3.0 (Waters).

For the continuous deuterium labelling (HDX), TcPOP (48 μM) was diluted 1:50 in a deuterated buffer at 20 mM HEPES, 150 mM NaCl (96.5% D_2_O fraction, pH_read_ 7.0) and the exchange reaction was conducted for 2 sec, 10 sec, 100 sec, 1,000 sec, 10,000 sec and 18 hrs at room temperature. The exchange reactions were quenched by a 1:1 dilution into ice-cold 100 mM phosphate buffer containing 3 M urea and 70 mM tris(2-carboxyethyl) phosphine (TCEP) (final pH_read_ 2.3). Samples were held on ice for 30 sec, snap-frozen in liquid nitrogen and kept frozen at –80°C until LC-MS analysis. A maximally labelled sample (MaxD) was produced by labelling TcPOP with 3 M fully deuterated urea in D_2_O and 2.5 mM TCEP, resulting in a final deuterium content as for the other labelled samples. The maximally labelled sample was quenched after 16 hrs by 1:1 v/v dilution into ice-cold 100 mM phosphate buffer (final pH_read_ 2.3), held for 30 sec on ice, snap-frozen in liquid nitrogen and kept frozen at −80°C until LC-MS analysis. A pulse labelling experiment (two technical replicates) was conducted to assess the protein stability and rule out irreversible unfolding of the protein over time. An aliquot of TcPOP was quickly unthawed and immediately labelled for 10 sec, then exposed to room temperature on the bench for 10,000 sec (as under conditions of continuous labelling) and labelled again for 10 sec. Frozen protein samples were quickly thawed and injected into an Acquity UPLC M-Class System with HDX Technology (Waters). The protein was on-line digested at 20°C into a home-made pepsin column and trapped/desalted with solvent A (0.23% formic acid in water, pH 2.5) for 3 min at 200 μL/min and at 1 °C through an Acquity BEH C18 VanGuard pre-column (1.7 μm, 2.1 mm × 5 mm, Waters). Peptides were eluted into an Acquity UPLC BEH C18 analytical column (1.7 μm, 2.1 mm × 100 mm, Waters) with a 7 min-linear gradient raising from 8 to 35% of solvent B (0.23% formic acid in acetonitrile) at a flow rate of 40 μL/min and at 1 °C. Then, peptides went through electrospray ionization in positive mode and underwent MS analysis with ion mobility separation.

Peptide level deuterium uptake was calculated with DynamX 3.0 (Waters) and data visually inspected and curated. Selected peptides exhibiting evident bimodal HDX behaviour (a clear sign of correlated exchange) were analysed by HX-Express3^53^ (HX-Express Software). Binomial fitting was applied with optimized fits for the number of amides, and undeuterated mass envelopes were calculated from the peptide sequence and fitted into the experimental undeuterated mass envelope to check the agreement. Subsequently, bimodal deconvolution (double binomial) was enabled for mass envelopes flagged as bimodal. The relative deuterium uptake (Da) and population fraction of both low- and high-mass envelopes were calculated. The fraction of the high-mass population (*y*) over the time points studied (*x*) was subjected to single or double exponential fitting by HX-Express3 to calculate the kinetic parameter Kop (rate of opening), according to the following exponential function:

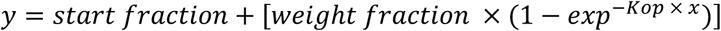

To determine the number of backbone amide hydrogens undergoing correlated exchange (#NHs) for the individual peptides, the time point showing maximal difference in HDX between the low- and high-mass population (Max ΔHDX) was identified. Then, the Max ΔHDX value was normalized by the peptide MaxD uptake and corrected for the number of exchangeable amides (N) and D_2_O fraction (= 0.9457), according to the following equation, as previously described^54,55^:

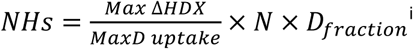

## Results

### Expression and purification of TcPOP and TcPOP homologues

Prolyl oligopeptidases (POPs) belong to the large hydrolase family, which is highly conserved in eukaryotes. To explore the evolutionary and functional similarities, we compared *T. cruzi POP* (TcPOP) to two close parasite homologues, namely those in *T. brucei* (TbPOP), *L. infantum* (LiPOP), and to the *Homo sapiens* homologue (HuPOP). Phylogenetic analysis and sequence conservation mapping on TcPOP highlighted that conserved and variable regions are located in both the catalytic and non-catalytic domains (Fig.S1a). Notably, the AlphaFold3 modelling of collagen binding to TcPOP corroborated previous docking studies with the substrate-binding site at the interface of the two domains (Video S1). This is a region of high variability, as indicated by the phylogenetic analysis (Fig.S1b). ^22^ TcPOP was expressed in *E. coli* using bacterial fermentation, overcoming the challenge of extremely low expression yields in batch culture (∼ 0.05 mg/L) (Fig. 1a). In contrast, expression of the other homologue proteins led to higher protein yields (0.8-2.0 mg/mL) (Fig.S2a). Following purification by size-exclusion chromatography (SEC) (Fig. S2b), all POPs exhibited enzymatic activity (Fig. S3a–c), thermostability, as determined by differential scanning fluorimetry (DSF) (Fig. S3d), and comparable end-point kinetics consistent with previously reported values for this enzyme family^56^. SEC-MALS analysis confirmed that all proteins were monomeric and monodisperse in solution (Fig.1b) and predicted the expected mass within the experimental error of the technique (TcPOP: 73.4 (±2.1%) kDa, TbPOP: 73.7 (±1.6%) kDa, LiPOP: 75.7 (±3.5%) kDa, HuPOP: 85.3 (±1.71%) kDa) (Fig. S4a). AlphaFold3 modelling of TcPOP predicted a structure comprising an α/β-hydrolase domain housing the catalytic triad Ser548-Asp631-His667, and a seven-bladed beta-propeller non-catalytic domain (Fig. S5), consistent with the previously determined structure of porcine POP (PDB code 1QFM)^22^.

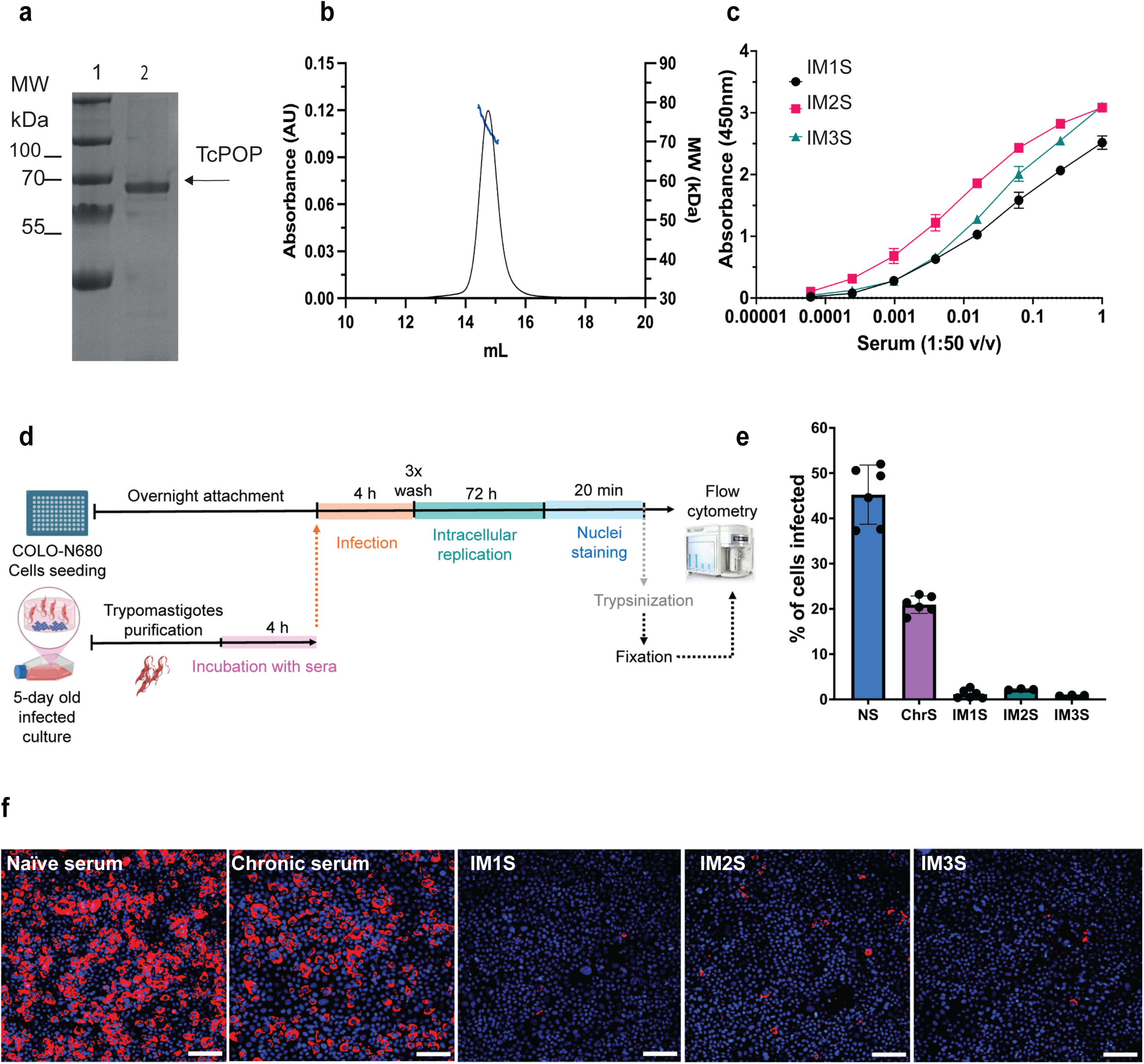

### Anti-TcPOP polyclonal response is cross-reactive

To assess the cross-reactivity of antibodies raised against TcPOP, mice were immunized with the recombinant enzyme. Homologous proteins with high amino acid identity from *L. infantum* (LiPOP, 63%), *T. brucei* (TbPOP, 73%), and *H. sapiens* (HuPOP, 43%) were expressed and purified to evaluate potential cross-reactivity, driven by the likelihood of shared epitopes (Fig.S1a). Three mice were immunised with TcPOP and polyclonal response analysed by ELISA (Fig.1c). The corresponding TcPOP-specific sera were tested against all four POPs. Cross-reactivity was detected to TcPOP, TbPOP, and LiPOP, but not for HuPOP at serially diluted serum concentrations. These findings suggest potential epitope conservation across parasite species, but not with HuPOP (Fig. S6a).

### Anti-TcPOP polyclonal sera block parasite invasion

To evaluate the neutralizing properties of anti-TcPOP polyclonal sera, we adapted our trypomastigote drug screening procedure ^57^ to a cell-based invasion assay (Fig.1d). Polyclonal antisera from all three mice demonstrated significant neutralizing activity, greatly reducing trypomastigote invasion of COLO-N680 cells compared to the naïve serum and serum derived from mice chronically infected with *T. cruzi.* (Fig. SS7a). Overall, we found that all three independently derived sera achieved a >95% neutralising effect, an impact that approached 99% for the serum from mice 1 (IM1S (Fig.1e-f), as measured by flow cytometry 72 hrs post infection (Supplementary Fig. S8a-b). The potency of IM1S was maintained even when reduced to 2.5% (Fig. 2a-b and Fig. S7b).

Immunofluorescence analysis of freshly extracted and fixed parasites revealed that the anti-TcPOP antisera bound in the endocytic/exocytic pathway of trypomastigotes (Fig. 2c, Supplementary Fig. S9b and Video S2). This pattern differed from antiserum from chronically infected mice, which showed the expected dispersed binding on the parasite surface, associated with a non-protective response (Fig. 2c). To understand the dynamics of IMIS binding, we performed a real-time assay which revealed that antibody attachment led to parasite lysis within minutes. To dissect this effect, we monitored the process by reducing the amount of serum and decreasing the temperature from 37°C to 30°C. (Fig. 2d, Supplementary Fig. S9a). Even under these constrained conditions, parasites became swollen after 5 mins exposure, followed by intracellular leakage and parasite lysis within 15 mins. (Fig. 2e, Supplementary Fig. S9c). To confirm this lytic effect, we incorporated a competition assay, including foetal bovine serum (FBS) at the same concentration as the IM1S. At the early timepoints, parasite integrity was preserved, and the majority of binding was restricted to the parasite surface. (Supplementary Fig. S9d).

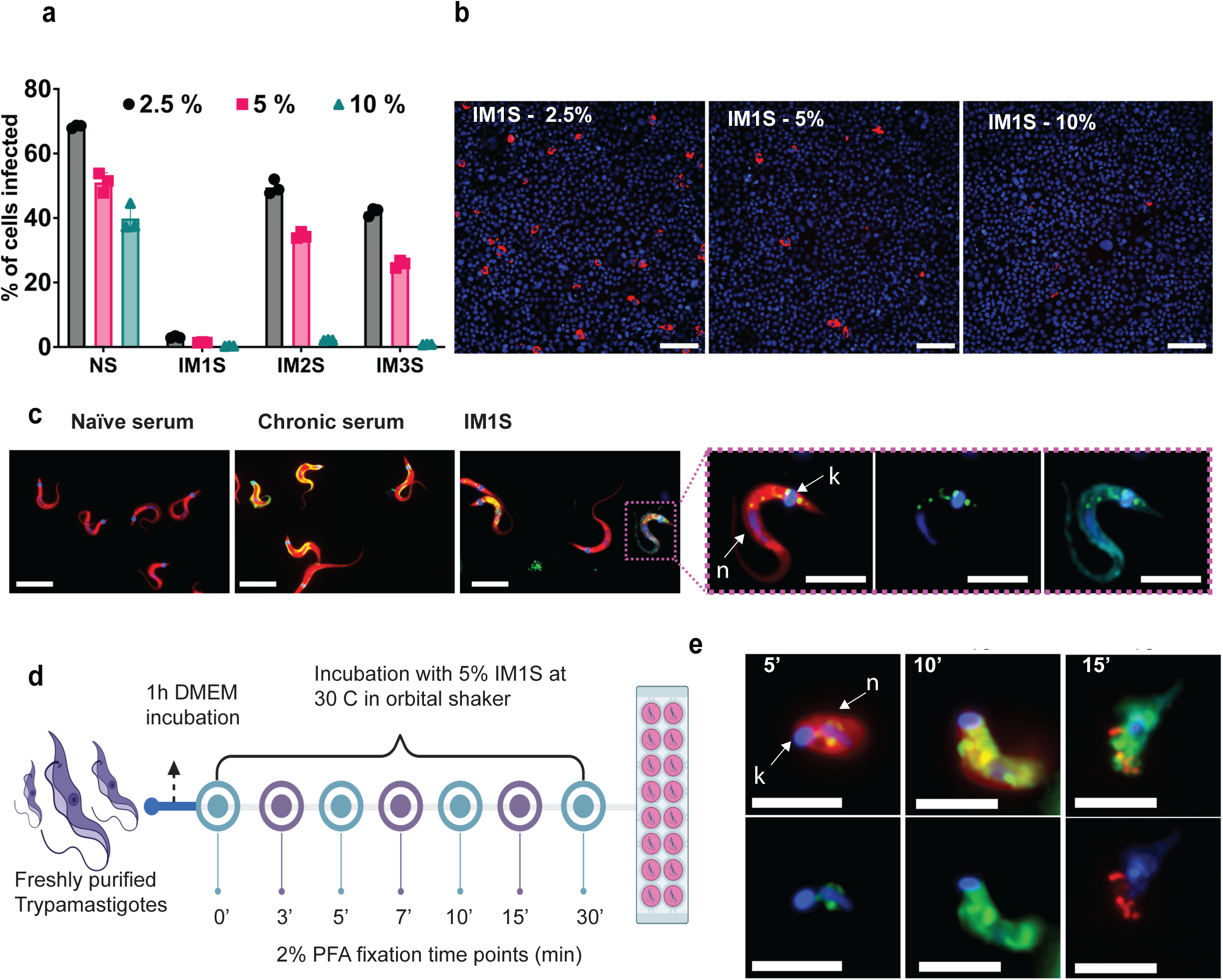

Additionally, we investigated binding to amastigotes, the intracellular form of the parasite. This life-cycle stage is also infectious, for example, when are released as host cells rupture following trypomastigote egress. The amastigote binding profile also highlighted the endocytic/exocytic pathway (Supplementary Fig. S9b). However, in live-binding assays, we also observed that surface attachment of anti-TcPOP, located at the apical end of the amastigote (Supplementary Fig. S9c-d). Finally, we determined binding in epimastigotes, the replicative insect form of the parasite The main location was at the anterior end, adjacent to the cytostome-cytopharynx complex, an additional secretory portal found in this form of the parasite. (Supplementary Fig. S9c-d). Overall, the binding studies reveal that TcPOP antisera is associated predominantly with the secretory pathway in all forms of the parasite.

### Anti-TcPOP monoclonal response

Three anti-TcPOP monoclonal antibodies (mAbs) were generated from the spleen of the mouse with the highest polyclonal serum response (Fig.3a) and screened by ELISA for specificity against TcPOP (Fig.3b) and homologue POPs (Fig.S6b). TcPOP antibody titres were positive at 1 µg/ml for mAb1 and at 0.1 µg/ml for mAbs 2 and 3 (Fig.3b), while no binding was observed against LiPOP, TbPOP and HuPOP. HRP-conjugation of anti-TcPOP mAbs revealed the presence of both linear and conformational epitopes, as confirmed by Western blot analysis (Fig. 3c).

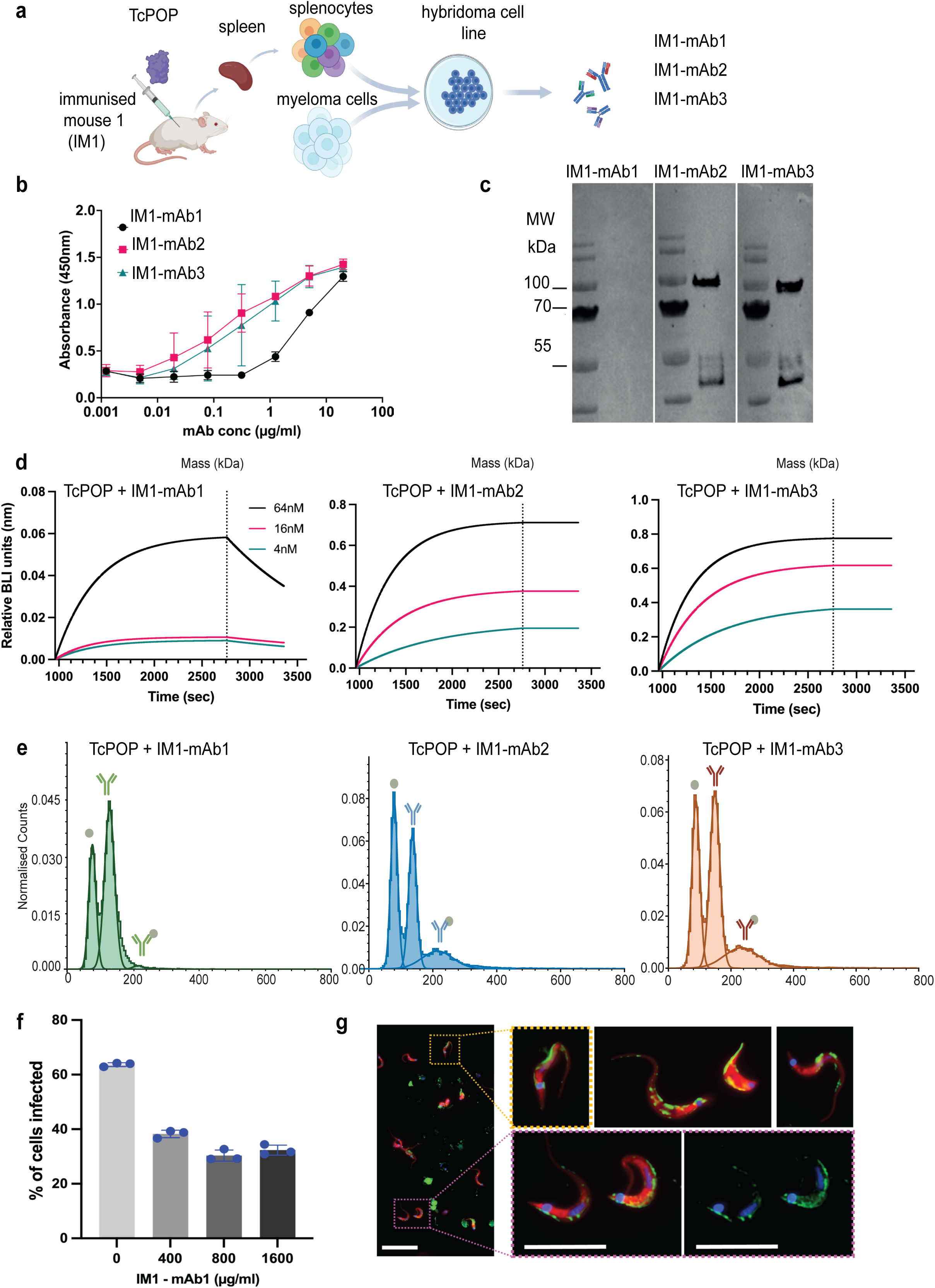

The BLI-based binding profile for immobilised anti-TcPOP mAbs against TcPOP showed higher binding affinity to TcPOP for anti-TcPOP mAb2 (K_D_ <1 pM, K_a_ (1/Ms) 2.3 x 10^5^, K_diss_ (1/s) < 10^-7^) and anti-TcPOP mAb3 (K_D_ <1 pM, K_a_ (1/Ms) 3.1 x 10^5^, K_diss_ (1/s) < 10^-7^). Instead, anti-TcPOP mAb1 exhibited comparatively weak binding (K_D_ < 3.2 nM, K_a_ (1/Ms) 7.1 x 10^4^, K_diss_ (1/s) 6.2 x 10^-4^) (Fig. 3d), as well as association and dissociation parameters. The best two mAbs, mAb2 and mAb3, were used to perform a sequential affinity binding with association and dissociation length of 1800 sec and 600 sec respectively. Both mAbs showed detectable association and dissociation (K_D_ <1 pM, K_a_ (1/Ms) >10^7^, K_diss_ (1/s) < 10^-7^) indicating competition for different epitopes (Fig.S11c).

Mass photometry measurements of TcPOP and the three individual anti-TcPOP mAbs, were 85 kDa and 151 kDa, respectively, with only a single peak present in each MP spectrum (Fig.S11a-b). Interaction studies revealed similar binding between TcPOP and both antibodies. A peak corresponding to 1:1 (mAb to TcPOP) was observed at 233 kDa with a relative abundance of 1-2% (TcPOP-mAb1), 17% (TcPOP-mAb2) and 19% (TcPOP-mAb3). No evidence of higher-order binding was found (Fig.3d).

### Anti-TcPOP monoclonal antibody neutralises parasite invasion

To further investigate the neutralising activity of the mAbs, we performed a cell-based infection assay incubating the parasites in 1000 µg/ml of each purified antibody in DMEM. Notably, mAb1 caused a significant reduction in parasite invasion, approaching 50% (Fig. 3f, Supplementary Fig. S10a). Additionally, we found that this neutralising effect was maintained regardless of the concentration assayed (Fig. 3f), and that instead of inducing a lytic effect, the effect was mediated by steric hinderance resulting from the binding to the surface of the trypomastigotes (Fig. 3g). The colocation assay showed a specific binding in a line along the flagellum (Supplementary Fig. S10b-c, Supplementary Video S3). Additionally, we observed that in amastigotes, the binding also associated with the short flagellum, that in these rounded forms of the parasite, is largely confined to the flagellar pocket (Supplementary Fig. S10b-c).

### Determination of cryo-EM structure of TcPOP in multiple conformations

#### TcPOP structure determination using single particle cryo-EM

We analysed the structure of TcPOP using single particle analysis (SPA) cryo-EM. 2D classification highlighted equal distribution of the particles in the micrographs (Fig.S12). The initial data collection comprised of 20,080 micrographs revealed severe preferential orientation^58^ resulting in poor map reconstructions (Fig.S13). Several approaches were attempted in cryo-EM grid preparation including changes in detergent, support films and glow discharge parameters. However, none led to significant improvement in the orientation distribution. We therefore tilted the grids to 30° and 35° to increase the number of views visualised in the electron micrographs (Fig.S13). This significantly improved the number of views, and the map produced was drastically improved with continuous density and minimal anisotropic features (Fig.S13). 3D classification revealed two distinct conformations of TcPOP i.e. open and closed (Fig.S13), indicating intrinsic conformational heterogeneity of the dataset. Closed and open conformation models obtained from AlphaFold and homology modelling (using PDB 3IUJ), respectively, were then docked into their respective CryoEM maps and refined (Figure 4a-b, Table 2). Local resolution was also Table 2 Cryo-EM statistics for closed and open conformation from real-space refinement in PHENIX^59^.

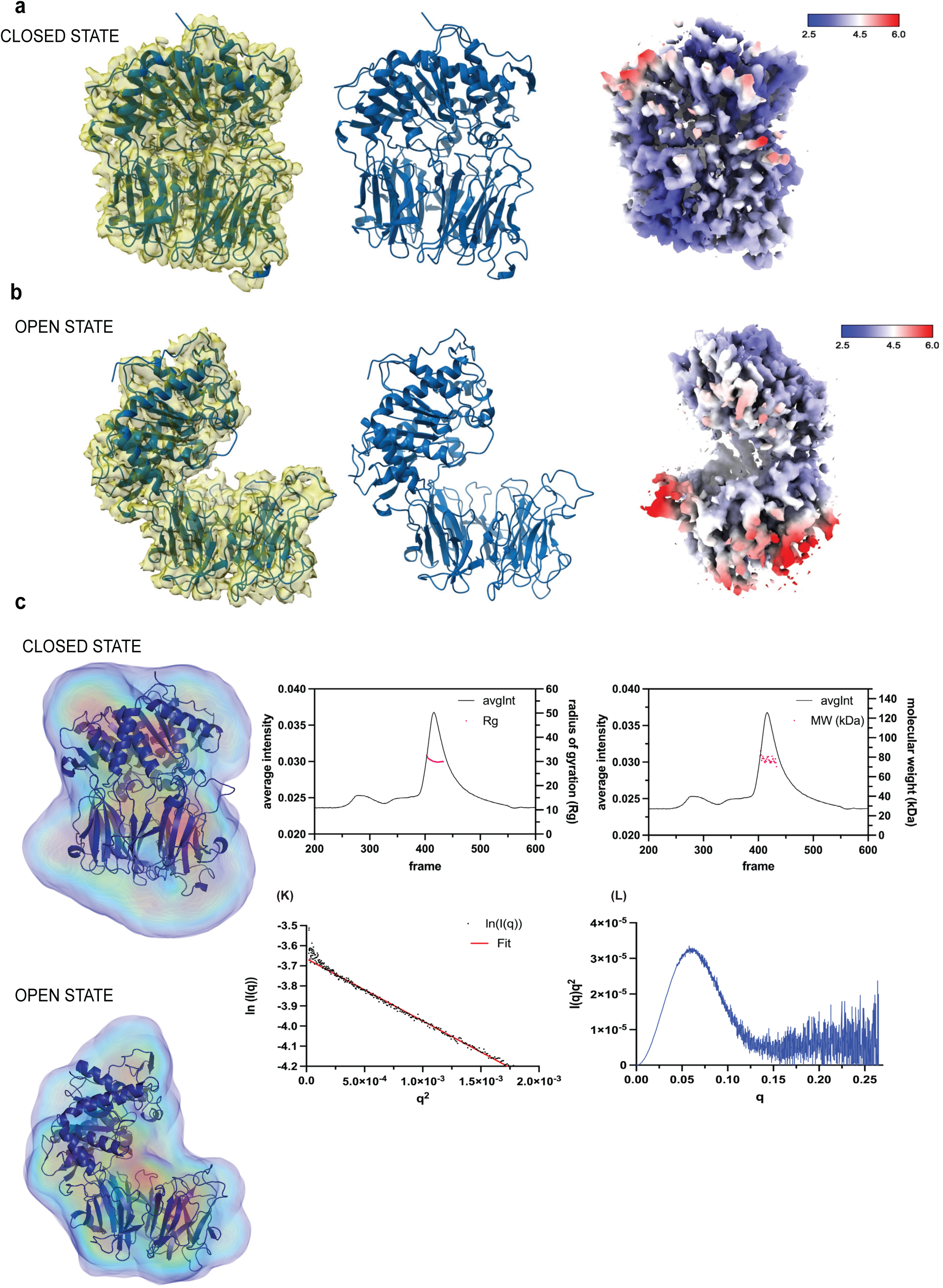

**Table 2.** Cryo-EM statistics for closed and open conformation from 610 real-space refinement in PHENIX^59^.

| **Refinement parameters** | **Closed conformation** | **Open conformation** |
| --- | --- | --- |
| PDB entry | 9HJI | 9HJJ |
| FSC (model) (masked) = 0.143 | 3.57 Å | 3.82 Å |
| Correlation Coefficient (masked) | 0.7 | 0.7 |
| Molprobity Score | 2.42 | 2.54 |
| Clash score | 12.95 | 14.79 |
| Ramachandran allowed | 100 % | 100% |
| Ramachandran favoured | 94.39% | 90.79 |
| Ramachandran outliers | 0% | 0% |
| Poor rotamers | 3.51% | 2.84% |
| Overall B factor (isotropic) | 70 | 90 |
| Bad bonds (RMSD) | 0% | 0% |
| Bad angles (RMSD) | 0.01% | 0.01% |

Our cryo-EM data confirm that TcPOP comprises two distinct domains: a catalytic α/β-hydrolase domain, which houses the active site featuring the catalytic triad, and a β-propeller domain, which likely regulates substrate access. The spatial arrangement of these domains reveals a cylindrical structure, with the β-propeller domain capping the catalytic α/β-hydrolase domain, as previously postulated^60^. This configuration suggests a gating mechanism controlling substrate entry to the active site, which may play a critical role in modulating enzymatic activity. These findings align with the proposed role of TcPOP in hydrolysing large substrates and facilitating parasite invasion into host cells.

#### SAXS and mass photometry analysis

SEC-SAXS profiles on purified TcPOP sample indicates no aggregation and globular conformation leading to an estimated molecular weight of 73.4 kDa ^61^, based on Kratky plot and Guinier analysis Open and closed conformations were also de-convoluted and the corresponding cryo-EM models were docked into the molecular envelopes (Fig.4c). Static SAXS data were also collected in a range concentrations, including zero-concentration, to determine the optimal level for SAXS data analysis and for data processing (Fig.S15). Mass photometry also allowed identification of the best buffer conditions in term of ionic strength and pH values (Fig.S11a). This information was exploited for cryo-EM grid preparation to increase sample homogeneity and also for buffer optimisation to elucidate TcPOP-mAbs interaction in BLI.

#### MD simulation of TcPOP

To understand the intrinsic dynamicity of TcPOP, molecular dynamics simulations were performed using the AlphaFold model from Uniprot (Q71MD6) as the starting point for 500 nsec simulations, which showed notable transitions in Root Mean Square Deviation (RMSD) around 290 nsec, while Root Mean Square Fluctuation (RMSF) (Fig.5a) and Solvent Accessible Surface Area (SASA) and radius of gyration (Rg) (Fig. S16) underwent complementary contraction for the first 300 nsec, followed by significant expansion up to 500 nsec, indicating substantial conformational changes. RMSF analysis identified residues 192-198 and 306-333 as highly flexible. The Gibbs free energy landscape analysis identified four significant minima basins, providing insights into local and global minima in a 2D and 3D projection of FEL, as described previously^62^ (Fig.5a).

#### Plasmonic Optical nanotweezers confirms TcPOP dynamicity in solution

Aperture-based plasmonic nano-tweezers revealed the conformational dynamics of single, unmodified TcPOP. TcPOP trapping events were detected by changes in transmission levels, filtered at a cut-off frequency of 1 kHz, whilst time traces of *ΔT*/*T_0_* of TcPOP trapped in the absence of substrate AMC (blue and purple, Fig.S17), and with 100 µM AMC substrate introduced to the trapping site. Addition of substrate at 100 µM concentration led to larger fluctuations in transmission, indicating TcPOP exhibiting greater dynamic fluctuations during enzymatic cycles than in its apo-state. However, in the absence of a substrate, distinct signal fluctuations were observed above the background of Brownian motions in the trapping well, suggesting free transition of multiple conformations of TcPOP in solution. This is further corroborated by distinct peaks in the probability density functions (PDFs) (Materials and Methods).

#### Structural dynamic analysis of TcPOP by HDX-MS

To gain insights on the kinetics of transition between the closed and open conformation in TcPOP, we applied hydrogen-deuterium exchange (HDX) mass spectrometry (MS) (Table S1 in HDX Supplementary data). HDX-MS enables the monitoring of the structural dynamics of proteins in solution, based on the rate at which individual backbone amide hydrogens exchange with deuterium atoms^63^. The two domains of TcPOP proved to be similarly dynamic, both containing regions of rapid, medium, or fast HDX (Fig. 5b, and S19 and S20). Remarkably, peptides of several helices and loops of the α/β-hydrolase domain and of a single loop of β-propeller domain (Fig. S18) exhibited bimodal isotopic envelopes with resolved high- and low-mass distributions, whose intensities interconvert over time: a clear sign of EX1 and EXx kinetics. This type of HDX behaviour is generally seen for groups of amides undergoing cooperative unfolding events (correlated exchange) and populating an open state for a sizeable time frame^63,64^. By pulse deuterium labelling, we established that those segments are not irreversibly unfolding over time and that TcPOP is in a dynamic equilibrium under our HDX conditions. Therefore, every opening event (determined by the rate constant k_op_) is reversible (Fig. S21). The bimodal isotopic envelopes were fitted^65^ to determine the number of amides involved in correlated exchange, the kinetic parameters k_op_, and HDX half-life of the closed state (Figs. S22a-j and S23, and Table S2 in HDX Supplementary data), which report on the relative rates at which individual helices/loops of TcPOP transition to the open conformation. We localized the correlated exchange at the level of the amides indicated in Fig. 5c. We observed that the loop 191-209 (β-propeller domain, at the interface with the α/β-hydrolase domain) and the helix 495-521 and adjacent loop 465-479 (α/β-hydrolase domain) manifest a starting open state of approximately 40% of the population, indicating that these specific segments adopt two initial non-interconverting distinct conformations, and subsequently very fast transition to the open state (average half-life <1,000 min at 23 °C). All other regions (helix and loop spanning residues 16-48, and helix 545-608 of the α/β-hydrolase domain) have an initial open state that approximates to 0%, with segment 16-48 interconverting very slowly to the open state (average half-life >2,000 min at 23 °C), and segment 545-608 interconverting at medium rate (average half-life between 1,000 and 2,000 min at 23 °C) (Fig. 5c-d and S23). Taken together, our HDX-MS data corroborate with the existence of the TcPOP open and closed conformations that could be resolved in cryo-EM, and with the presence of intermediate conformations that arise from the switching of selected helices and loops at different rates.

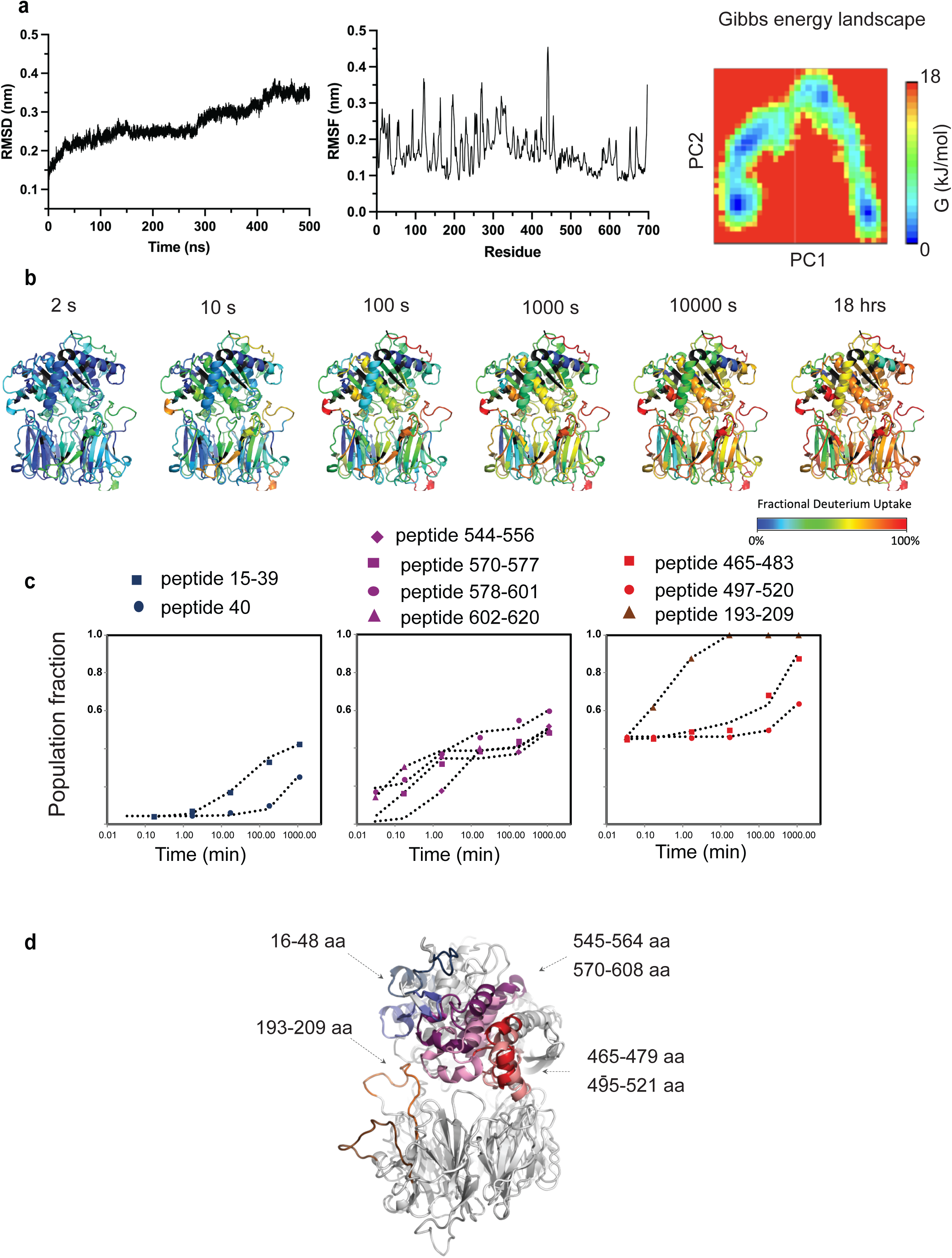

## Discussion

Structural data on *T. cruzi* antigens are desperately needed to guide vaccine development and design of diagnostic solutions. Such data are invaluable to initiate structure-based immunogen design strategies which focus the immune response towards key conserved epitopes^66^ and to identify regions within the target protein suitable for drug design. TcPOP is one of the leading validated candidates for a Chagas disease vaccine, given its high level of sequence conservation across the six genotypes (or DTUs) of the *T.cruzi* species.

The three-dimensional structure of TcPOP remained elusive for almost two decades since a comparative homology model was proposed in 2005^22^. Several attempt to crystalise parasite POPs in our lab, including TcPOP, LiPOP, TbPOP and HuPOP, did not succeed, despite the usage of starting protein concentrations up to 500 µM for crystallisation screening (equivalent to ∼ 40 mg/mL). Closed conformations for TcPOP homologues have been previously reported using protein crystallography from porcine POP (PDB code 1QFM), *Pyrococcus furiosus* (PDB code 5T88), or HuPOP with an irreversible inhibitor (PDB code 3DDU), to lock the enzyme in the closed state. However, to our knowledge, neither fully open nor fully closed conformations of a parasite POP in a single solution (with or without the aid of additives) have been determined previously. High solubility was a characteristic of not only TcPOP, but also of other POPs reported in this manuscript, and this is likely to be linked to their function of being secreted in the blood to perform degradation of the extracellular host matrix. In fact, TcPOP possess 33 Lys residues, out of which, 13 are located on the solvent accessible surface, a feature that likely prevents the crystal contacts required to promote crystallisation. Methods have been developed to reduce protein entropy by lysine methylation^67^ but we opted not to use this strategy, as this could have compromised antigenicity by altering potential key residues involved in epitope-paratope interaction.

Immunizing mice with recombinant TcPOP led to the production of polyclonal antibodies that showed cross-reactivity with *L. infantum* and *T. brucei* homologs, indicating conserved epitopes among these parasites. However, the lack of response to the human homolog led to the speculation that there might be a divergence in epitope conservation across species. This might be attributable to evolutionary pressure or possible immune suppression in the human host, although this remains to be established. Such observed cross-reactivity not only underscores the potential for developing cross-protective vaccines against related parasite species but also emphasizes the precision in targeting pathogen-specific epitopes, while minimizing cross-reactivity with human proteins for diagnostic applications. Here, we investigated the immune response in mice at both the polyclonal and monoclonal level to TcPOP. Polyclonal antibodies from all three mice exhibited striking neutralization effects, approaching 100%, and leading to rapid parasite lysis of the infectious trypomastigote stage. These findings consolidate TcPOP’s central role as a leading candidate for a Chagas disease vaccine. Additionally, the production and characterization of three monoclonal antibodies from the mouse with the highest polyclonal response identified a single mAb (IM1-mAb1) responsible for approximately 50% of the neutralization effects. Unexpectedly, this mAb, despite being neutralising, did not induce parasite lysis, suggesting the presence of an alternative neutralization mechanism, potentially involving hindrance, as indicated by cell localization experiments.

Interestingly, the IM1-mAb1was the weakest binder *in vitro* among the three isolated mAbs, exhibiting faster *kon* and *koff* rates, whilst the others exhibited much longer *koff* rates. This underscores the importance of testing all mAbs in a cell-invasion scenario that more closely reflects host-pathogen interactions, without excluding any based on *in vitro* studies alone, as interactions with the parasite may reveal neutralizing potential.

In our quest to determine the three-dimensional structure of TcPOP, the AlphaFold model of the closed conformation of TcPOP highlighted its dynamic nature, showing significant transitions in the Cα-backbone during the course of molecular dynamics (MD) simulations. Also, MD simulation suggested such transitions occur between the two domains of TcPOP exposing their catalytic site during enzyme activity. Importantly, the β-propeller domain contributed to a high degree of movement and instability, whereas, α/β hydrolase domain remained comparatively stable. Moreover, the Gibbs free energy landscape of TcPOP indicated scattered blue spots, representing four-to-six major local or global energy minima, and therefore, provided valuable insights into the presence of different metastable states (Fig. 5a). Coincidentally, the number of minima matched the number of classes in cryo-EM, possibly indicating that there might be a correlation between MD predictions and classes distribution, which will be further investigated for methods development.

We, therefore, decided to analyse the structure using cryo-EM, despite the challenge to date of resolving sub-80 kDa molecules, due to low image contrast^68^. To ensure the quality of samples before data collection, we implemented a quality control pipeline, which included SEC-SAXS in solution studies followed by assessment of the best buffer conditions using mass photometry to screen for the best conditions for monodispersion, therefore increasing the chances of success.

Our approach led to determination of the TcPOP structure to 3.6 Å and 3.8 Å, respectively, in closed and open conformation. This is one of the smallest cases, and at the highest resolution reported, in terms of resolving multiple enzyme conformations in a vitreous state by single-particle cryo-EM^69^. Two conformations of TcPOP could be easily identified in 3D classification spanning from fully closed to fully open with an overall motion of ∼ 22° between the two domains. The evident stability of the α/β hydrolase domain observed in both open and closed conformations, as well as in those from aligned cryo-EM maps, suggest possible region which are exposed to the immune system when the enzyme is secreted in the blood, and therefore likely to include potential epitopes.

To confirm that these experimentally observed conformations exist in solution, optical tweezer studies were performed in the presence and absence of substrate-mimicking peptide. Recent advances in plasmonic nano-tweezers allowed us to sample conformational fluctuations of single proteins in solution ^70^. Our data show that TcPOP clearly fluctuates between open and closed states independently from the presence of the substrate, confirming that the observed open and closed conformations occur in solution and are observed without the usage of any additive. However, the addition of the substrate-mimicking peptide affects the frequency at which the enzyme opens and closes.

By HDX-MS, we primarily observed that selected helices and loops of the α/β-hydrolase domain and a single loop of β-propeller domain (at the interface with the α/β-hydrolase domain) switch between the two conformations via cooperative unfolding/refolding events along the protein backbone, which are interpretable as long-lived perturbations of their secondary structure. This has been previously observed for other enzymes^64^. Such a kinetic regime allowed us to understand the structural elements of TcPOP switching between the closed and open conformation at different rates, which likely leads the protein to occupy a multitude of intermediate states between the fully closed and fully open conformation.

The intrinsic conformational heterogeneity in solution can pose great challenges to the structure determination by cryo-EM, as multiple conformations merge into a single conformation. This often leads to preferential orientation, eventually producing lower-resolution 3D reconstruction, which requires the collection of large number of micrographs especially for small molecules, often in the order of thousands. To overcome this barrier, we have analysed each of the distinct conformers with DynaMight. Here, we have successfully characterized the open and close conformation of the vaccine candidate TcPOP through a synergistic combination of *in silico*, structural, and in solution techniques. We propose expanding this methodology for regular investigations of small, secreted proteins of sub-80kDa size, especially when recalcitrant to crystallography approaches and where alternating conformations in solution, are suspected. The addition of tilted data collection can also be beneficial to alleviate preferential orientation, therefore reveal more structural details.

We envisage the possibility to extend this approach to other targets, allowing prediction of the number of 3D classes in vitreous states using cryo-EM data based on MD simulations. In addition, it will promote understanding of conformational heterogeneity at the early stages of cryo-EM data processing and therefore, potentially aid future software developments towards this goal. We provide experimental evidence on the distinct open and closed conformation, which will be invaluable to determine which regions of TcPOP (and potentially other members of prolyl oligopeptidase family) should be targeted to block the enzyme in either conformation, therefore aiding the development of novel and much needed anti-parasitic therapeutic agents.

Additionally, we characterised anti-TcPOP polyclonal response, which revealed striking neutralising properties with subsequent rapid lysis of trypomastigote. In addition, we isolated one monoclonal antibody, which is also neutralising, but via other means. This information highlights TcPOP as a priority vaccine target to be exploited for both, diagnostic and therapeutic solutions. Both are urgently needed to reduce the burden of Chagas disease. In the longer-term, our findings will allow a structure-guided development of a Chagas vaccine prototype, which is desperately needed in the fight against this major neglected disease.

## Contributions

S.B. expressed and purified proteins, characterised them biophysically and *in silico* and determined the cryo-EM structure. S.B. and C.L. prepared the samples for cryo-EM data acquisition. S.B., T.J.R., and E.H. processed and analysed cryo-EM data. S.B. and A.M.F. isolated and purified antibodies. L.B. immunised mice and performed FACS analysis. S.B., M.K. and W.B.S performed mass photometry experiments and data analysis. V.C. designed, performed and analysed HDX-MS. M.A. and C.Y. performed plasmonic optical tweezers studies. F.O. and J.M.K. performed infection and parasite binding experiments. I.C. conceptualise the experiments and provided funding. S.B. F.O. and I.C. prepared the manuscript and all authors contributed and commented on it.

## Acknowledgements

We would like to acknowledge: Dr Nathan Cowieson and Dr Katsuaki Inoue for assisting data collection at B21 beamline (Diamond Light Source), Dr David Staunton (University of Oxford) for data collection of SEC-MALS, Dr Lei Xu (NTU) and Prof. Rahmani Mohsen (NTU) for advice on data analysis of plasmonic optical tweezers experiments. Additionally, we would like to thank, Anu Itansanmi-Ogundayomi (Federal University of Technology Akure, Nigeria), Dr Jody Winter (NTU) and Dr Richard Cowan (University of Leicester). We would like to acknowledge Dr Maria Bassi and Kasper H Björnsson (University of Copenhagen) for advising in antibody methodologies, Dr David Owens (electron Bio-Imaging Centre, Oxon, UK) to provide advice in cryo-EM data quality assessment during the review process and Richard Atherton (LSHTM) for assistance with parasite cultures.

The project was funded by: Wellcome grant 204801/Z/16/Z (IC), Royal Society grant IES\R2\232167 (IC). SB is supported by the Nottingham Trent Doctoral School studentship and LB is funded by Novo Nordisk Foundation (NNF170C0026778). FO contribution was supported by Plan Propio of the University of Granada Research Stimulation grant (PP2023.PRI.I.14). We acknowledge The Midlands Regional CryoEM Facility at the Leicester Institute of Structural and Chemical Biology (LISCB), major funding from MRC (MC_PC_17136).

