## Supplementary Figures for "Cryo-EM led analysis of open and closed conformations of Chagas vaccine candidate TcPOP and its antibody response characterisation"

a

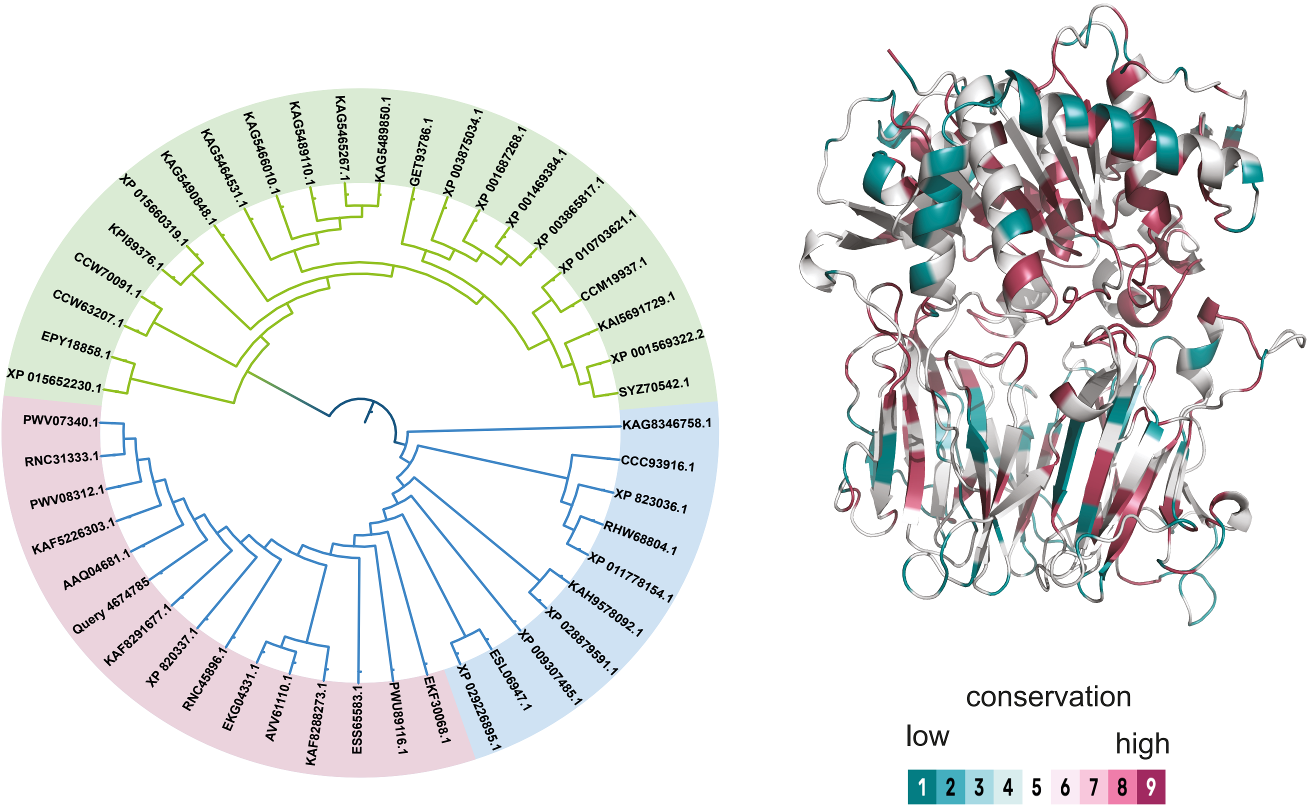
XP 015652230.1
EPY18858.1
CCW63207.1
CCW70091.1
KPI89376.1
XP 015660319.1
KAG5490848.1
KAG5464531.1
KAG5466010.1
KAG5489110.1
KAG5465267.1
KAG5489850.1
GET93786.1
XP 003875034.1
XP 001687268.1
XP 001469384.1
XP 003865817.1
XP 010703621.1
CCM19937.1
KAI5691729.1
XP 001569322.2
SYZ70542.1
KAG8346758.1
CCC93916.1
XP 823036.1
RHW68804.1
XP 011778154.1
KAH9578092.1
XP 028879591.1
XP 009307485.1
ESL06947.1
XP 029226895.1
EKF30068.1
PWU89116.1
ESS65583.1
KAF8288273.1
AVV61110.1
EKG04331.1
RNC45896.1
XP 820337.1
KAF8291677.1
Query 4674785
AAQ04681.1
KAF5226303.1
PWV08312.1
RNC31333.1
PWV07340.1
conservation
low
high
1 2 3 4 5 6 7 8 9

b

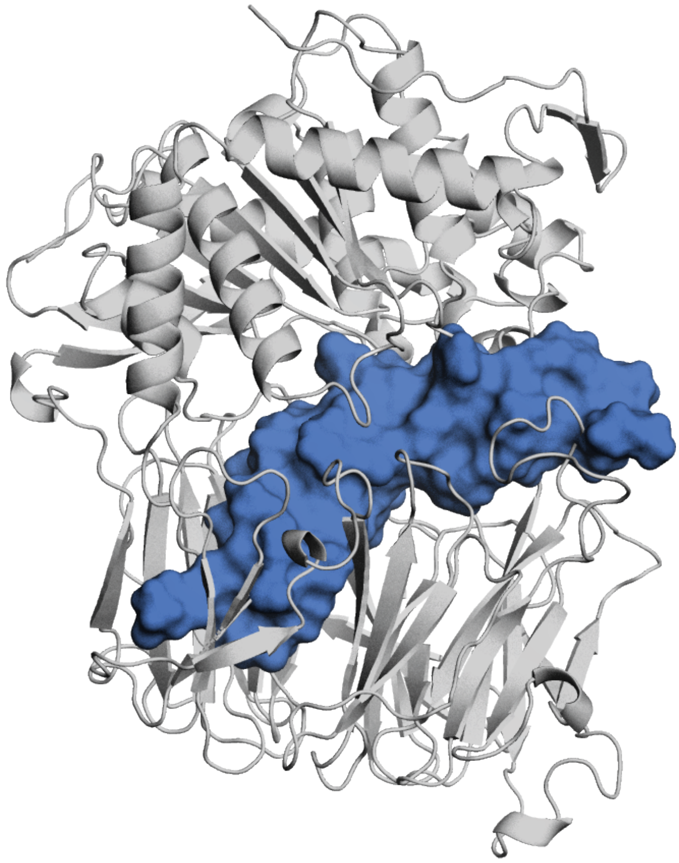

Figure S1

**a**

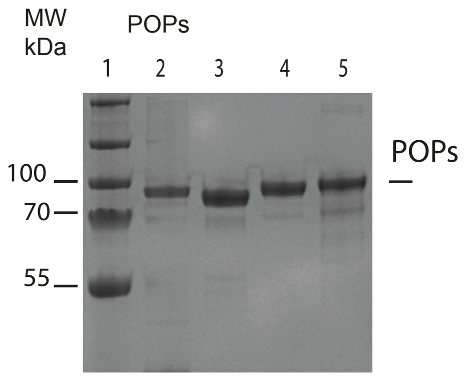
MW
kDa
POPs
1
2
3
4
5
100
70
55
POPs

**b**

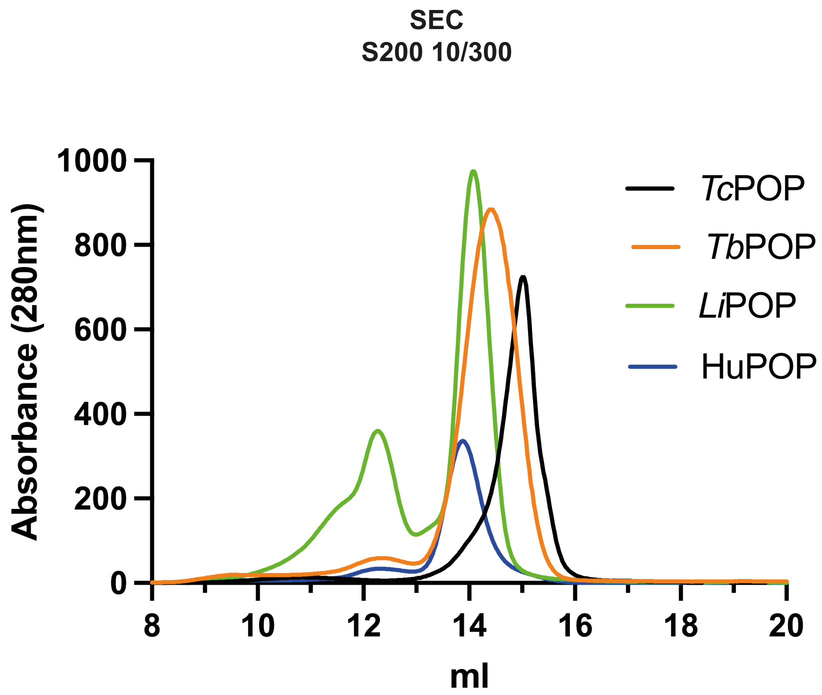
SEC
S200 10/300
1000
800
600
400
200
0
Absorbance (280nm)
8
10
12
14
16
18
20
ml
TcPOP
TbPOP
LiPOP
HuPOP

Figure S2

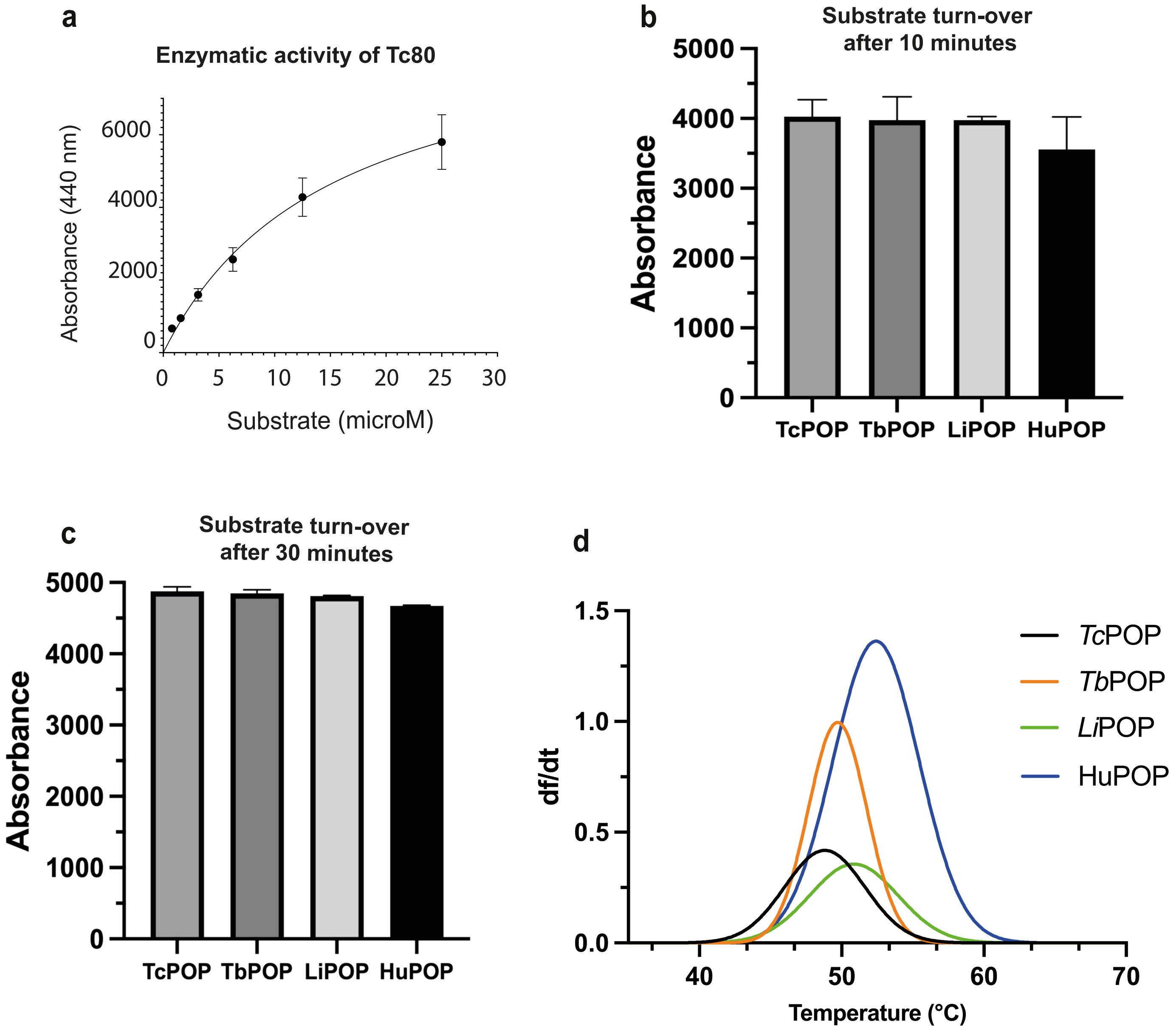

a
Enzymatic activity of Tc80
Absorbance (440 nm)
6000
4000
2000
0
0
5
10
15
20
25
30
Substrate (microM)
b
Substrate turn-over
after 10 minutes
Absorbance
5000
4000
3000
2000
1000
0
TcPOP
TbPOP
LiPOP
HuPOP
c
Substrate turn-over
after 30 minutes
Absorbance
5000
4000
3000
2000
1000
0
TcPOP
TbPOP
LiPOP
HuPOP
d
df/dt
1.5
1.0
0.5
0.0
40
50
60
70
Temperature (°C)
TcPOP
TbPOP
LiPOP
HuPOP


Figure S3

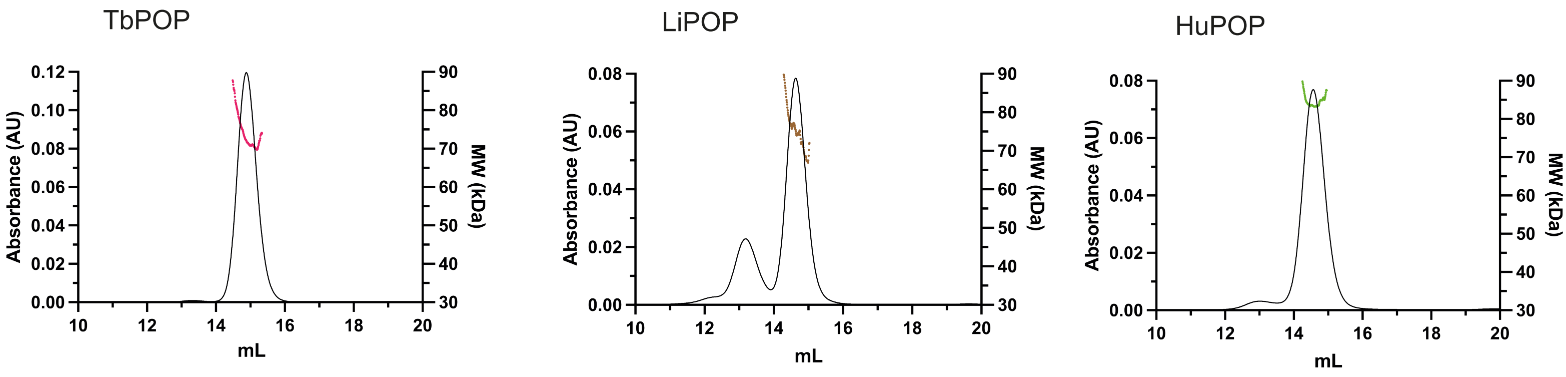

TbPOP
LiPOP
HuPOP
Absorbance (AU)
MW (kDa)
mL
0.00
0.02
0.04
0.06
0.08
0.10
0.12
30
40
50
60
70
80
90
10
12
14
16
18
20


Figure S4

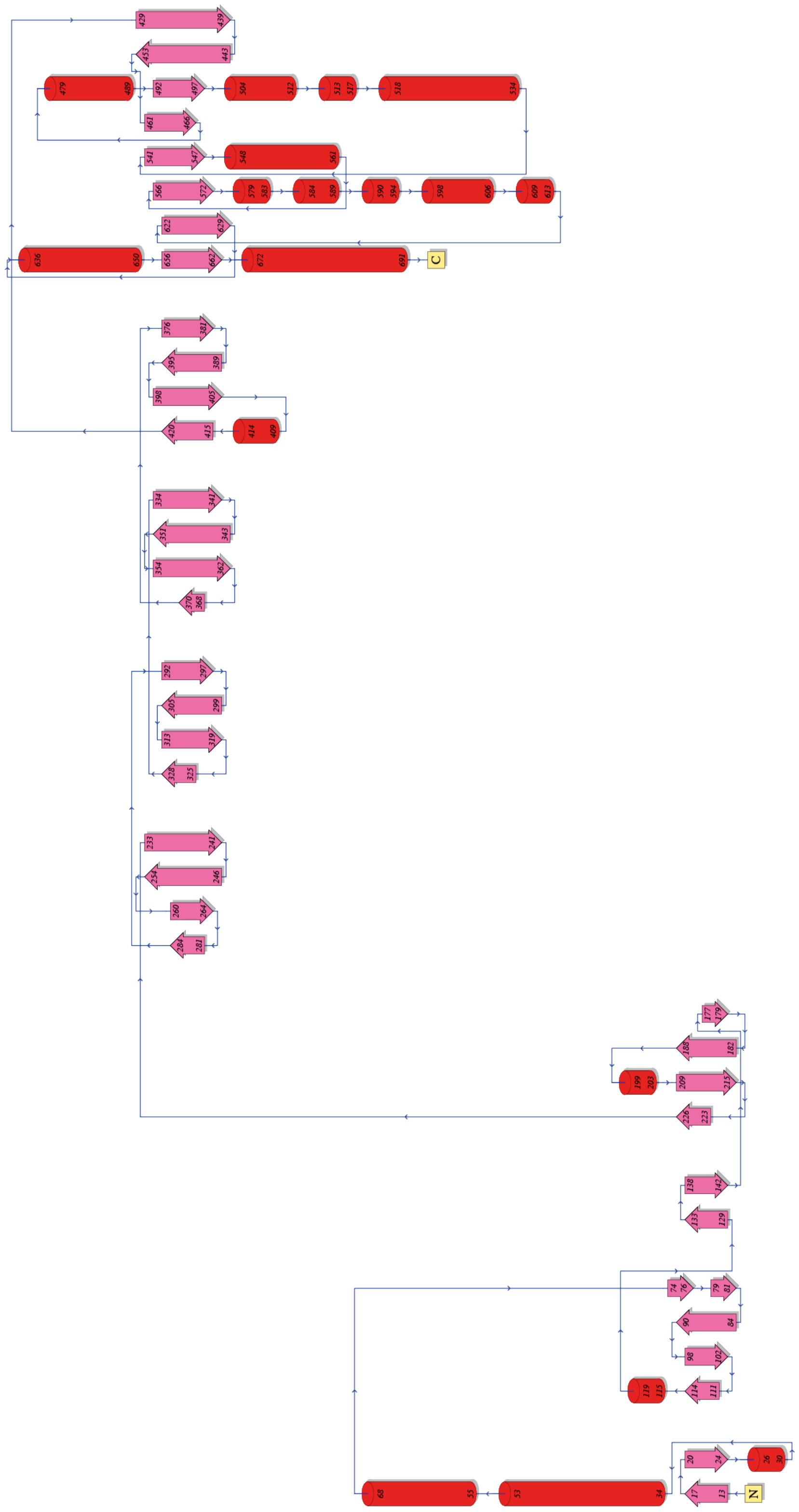


Figure S5

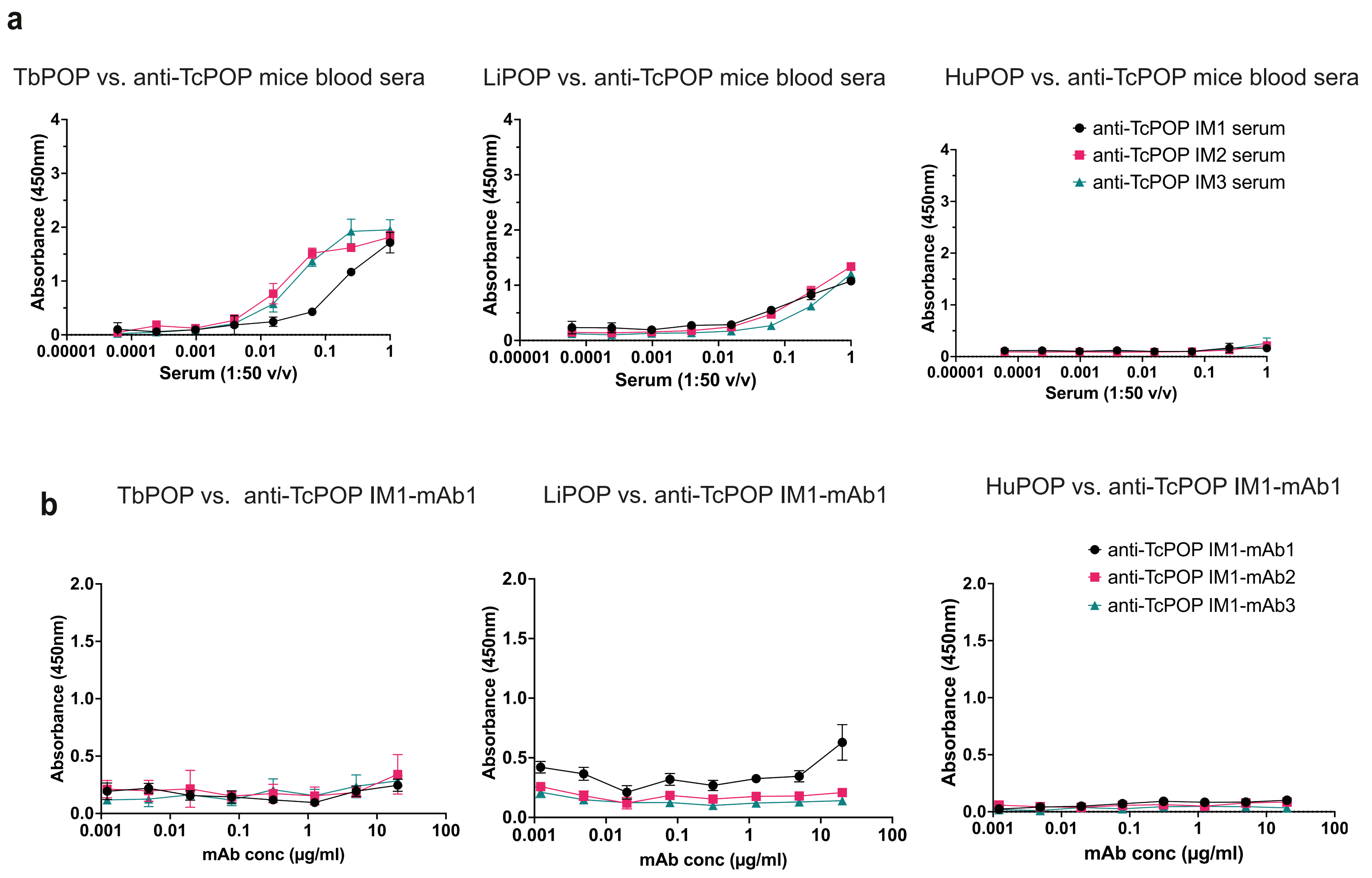
a
TbPOP vs. anti-TcPOP mice blood sera
LiPOP vs. anti-TcPOP mice blood sera
HuPOP vs. anti-TcPOP mice blood sera
anti-TcPOP IM1 serum
anti-TcPOP IM2 serum
anti-TcPOP IM3 serum
Absorbance (450nm)
Serum (1:50 v/v)
0.00001 0.0001 0.001 0.01 0.1 1
b
TbPOP vs. anti-TcPOP IM1-mAb1
LiPOP vs. anti-TcPOP IM1-mAb1
HuPOP vs. anti-TcPOP IM1-mAb1
anti-TcPOP IM1-mAb1
anti-TcPOP IM1-mAb2
anti-TcPOP IM1-mAb3
Absorbance (450nm)
mAb conc (μg/ml)
0.001 0.01 0.1 1 10 100

Figure S6

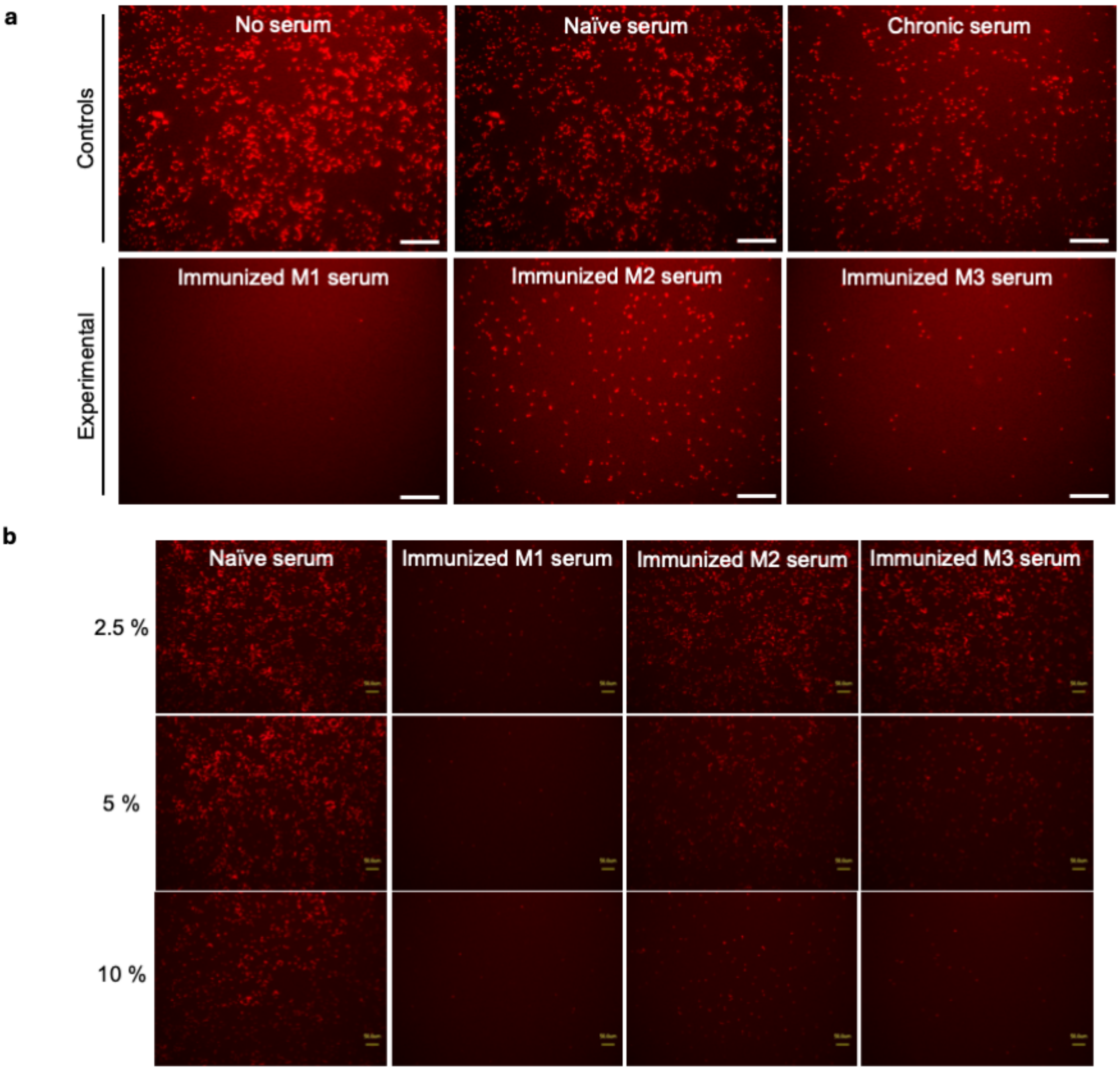
a
Controls
Experimental
No serum
Naïve serum
Chronic serum
Immunized M1 serum
Immunized M2 serum
Immunized M3 serum
b
Naïve serum
Immunized M1 serum
Immunized M2 serum
Immunized M3 serum
2.5 %
5 %
10 %

Figure S7

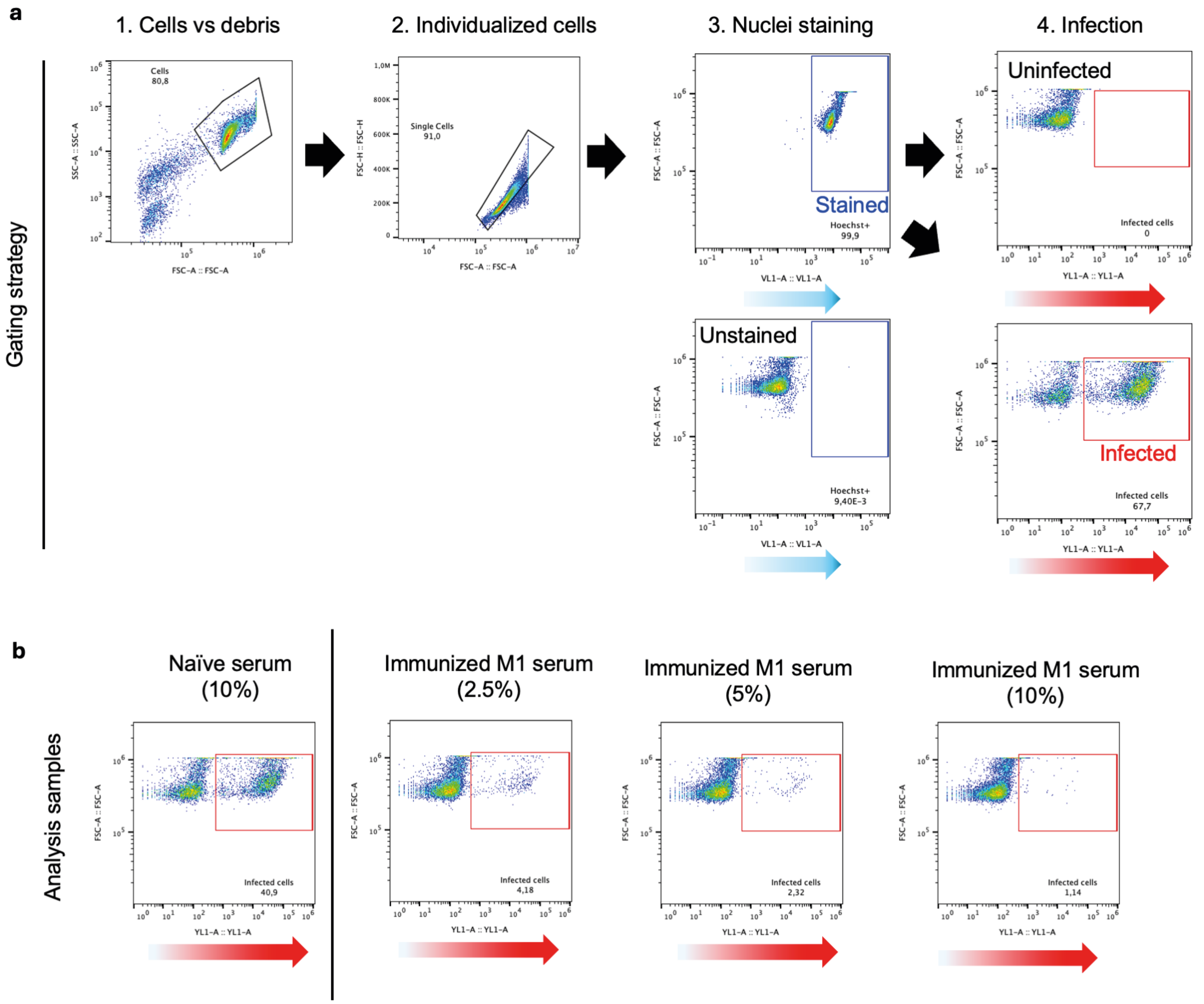
a
1. Cells vs debris
2. Individualized cells
3. Nuclei staining
4. Infection
Gating strategy
Cells
80,8
Single Cells
91,0
Stained
Hoechst+
99,9
Unstained
Hoechst+
9,40E-3
Uninfected
Infected cells
0
Infected
Infected cells
67,7
b
Analysis samples
Naïve serum
(10%)
Infected cells
40,9
Immunized M1 serum
(2.5%)
Infected cells
4,18
Immunized M1 serum
(5%)
Infected cells
2,32
Immunized M1 serum
(10%)
Infected cells
1,14

Figure S8

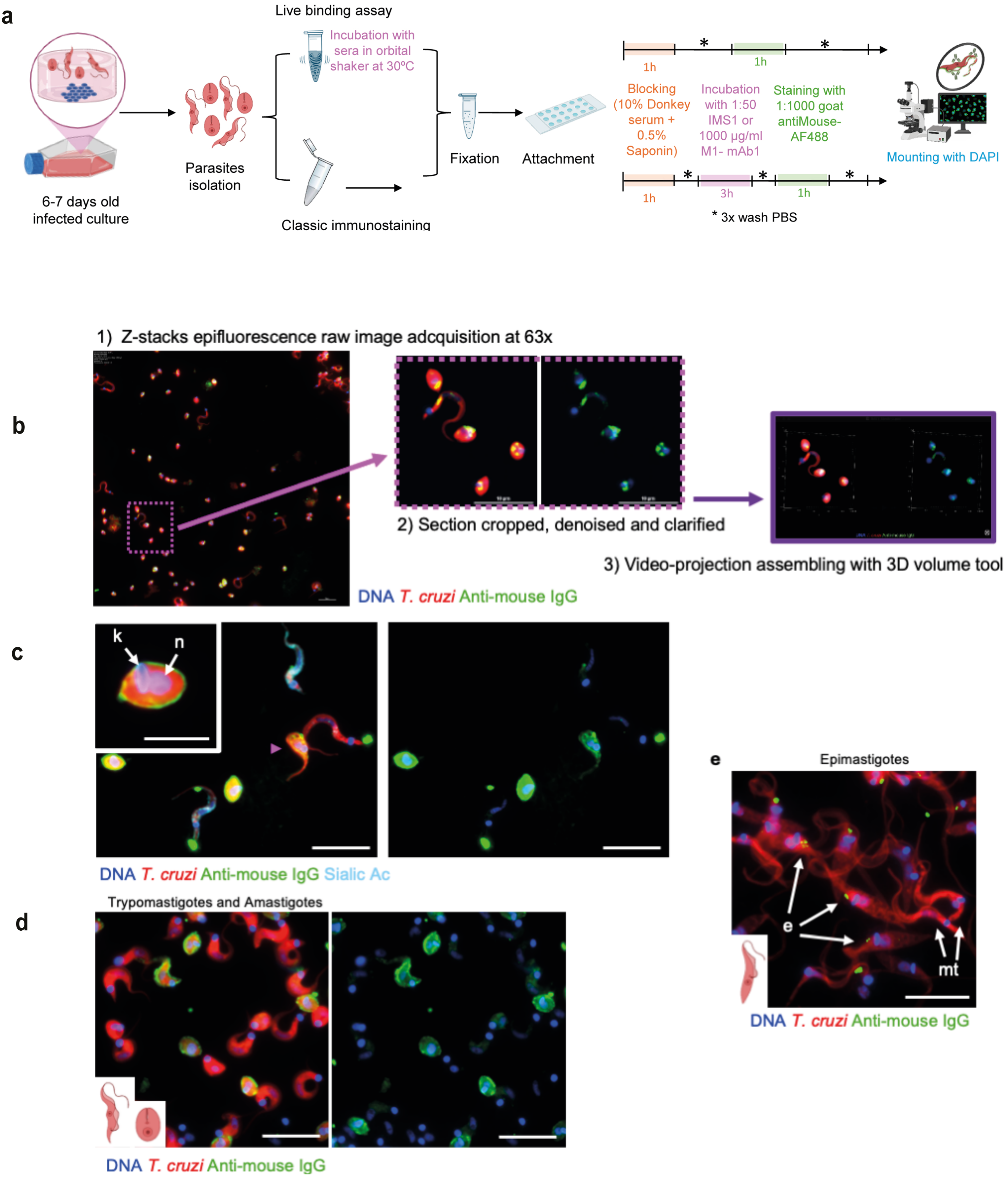

a
6-7 days old infected culture
Parasites isolation
Live binding assay
Incubation with sera in orbital shaker at 30ºC
Classic immunostaining
Fixation
Attachment
1h
1h
Blocking (10% Donkey serum + 0.5% Saponin)
Incubation with 1:50 IMS1 or 1000 µg/ml M1- mAb1
Staining with 1:1000 goat antiMouse-AF488
1h
3h
1h
* 3x wash PBS
Mounting with DAPI
b
1) Z-stacks epifluorescence raw image adcquisition at 63x
2) Section cropped, denoised and clarified
3) Video-projection assembling with 3D volume tool
DNA T. cruzi Anti-mouse IgG
c
k
n
DNA T. cruzi Anti-mouse IgG Sialic Ac
d
Trypomastigotes and Amastigotes
DNA T. cruzi Anti-mouse IgG
e
Epimastigotes
e
mt
DNA T. cruzi Anti-mouse IgG


Figure S9

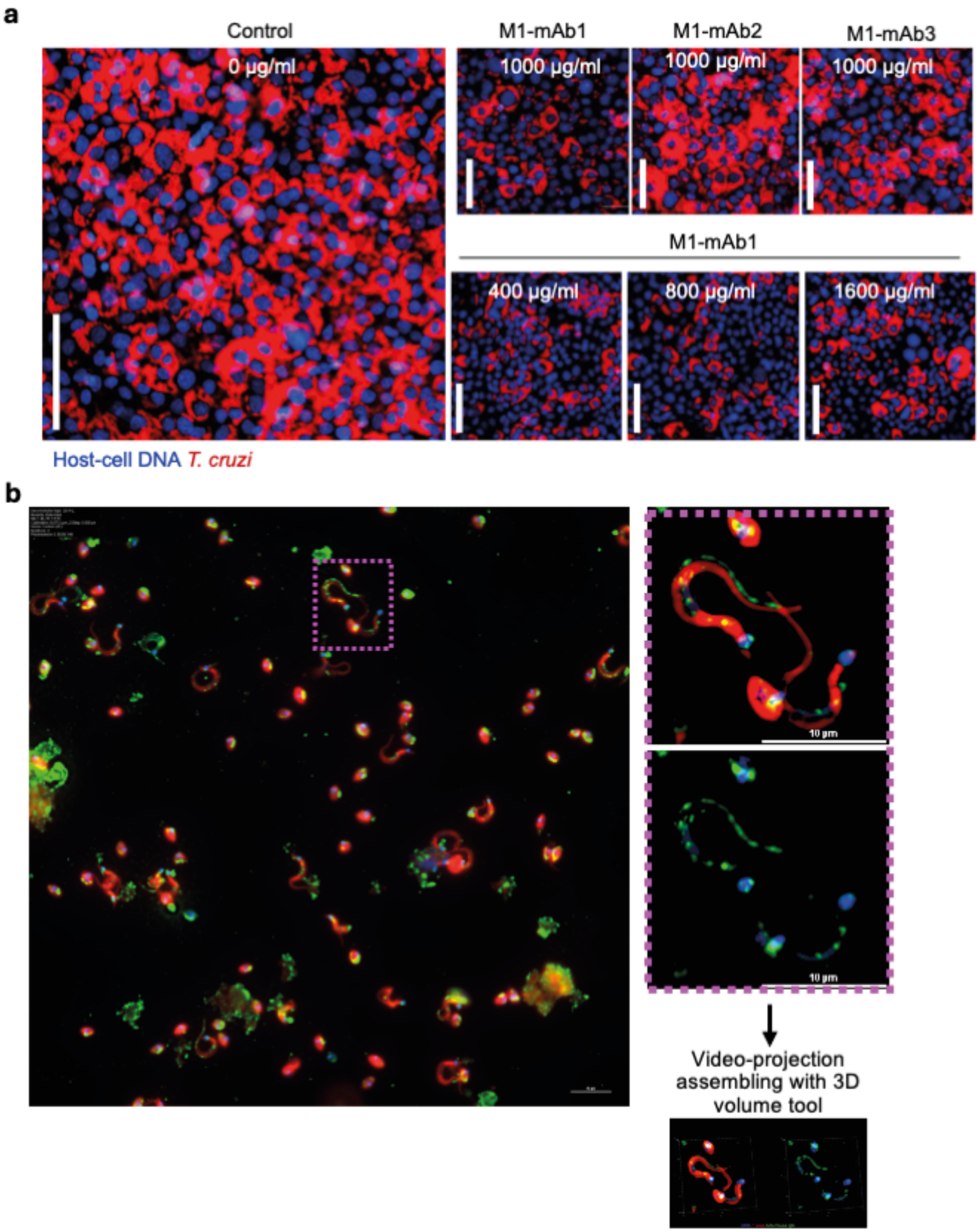
a
Control
0 µg/ml
M1-mAb1
1000 µg/ml
M1-mAb2
1000 µg/ml
M1-mAb3
1000 µg/ml
M1-mAb1
400 µg/ml
800 µg/ml
1600 µg/ml
Host-cell DNA T. cruzi
b
10 µm
10 µm
Video-projection assembling with 3D volume tool

Figure S10

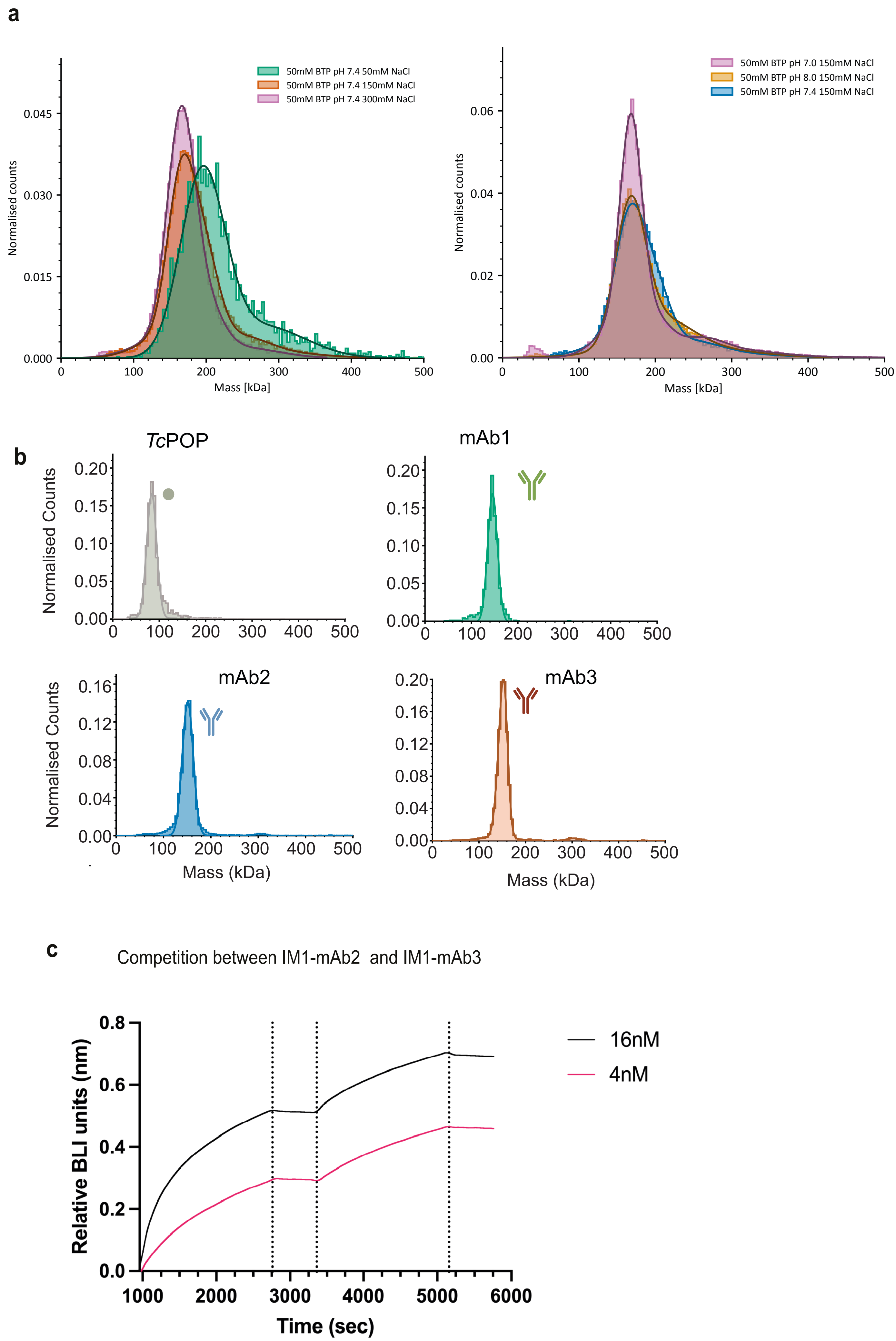
a
50mM BTP pH 7.4 50mM NaCl
50mM BTP pH 7.4 150mM NaCl
50mM BTP pH 7.4 300mM NaCl
Normalised counts
Mass [kDa]
50mM BTP pH 7.0 150mM NaCl
50mM BTP pH 8.0 150mM NaCl
50mM BTP pH 7.4 150mM NaCl
Normalised counts
Mass [kDa]
b
TcPOP
mAb1
mAb2
mAb3
Normalised Counts
Mass (kDa)
c
Competition between IM1-mAb2 and IM1-mAb3
16nM
4nM
Relative BLI units (nm)
Time (sec)

Figure S11

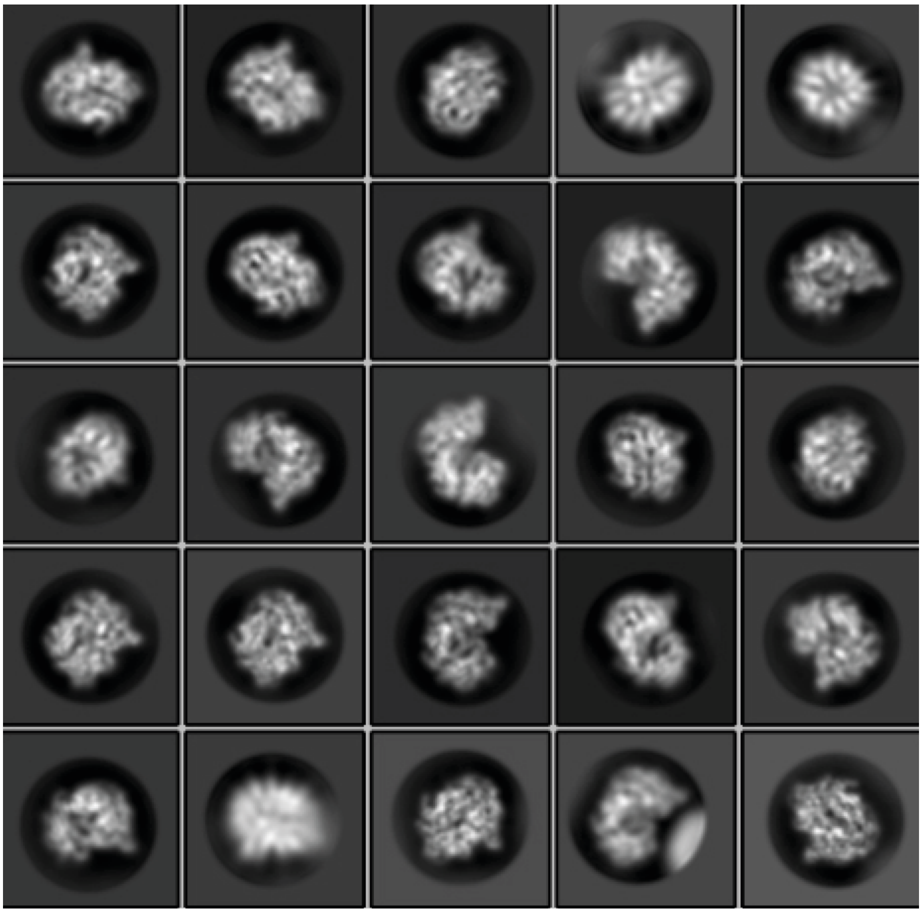

Figure S12

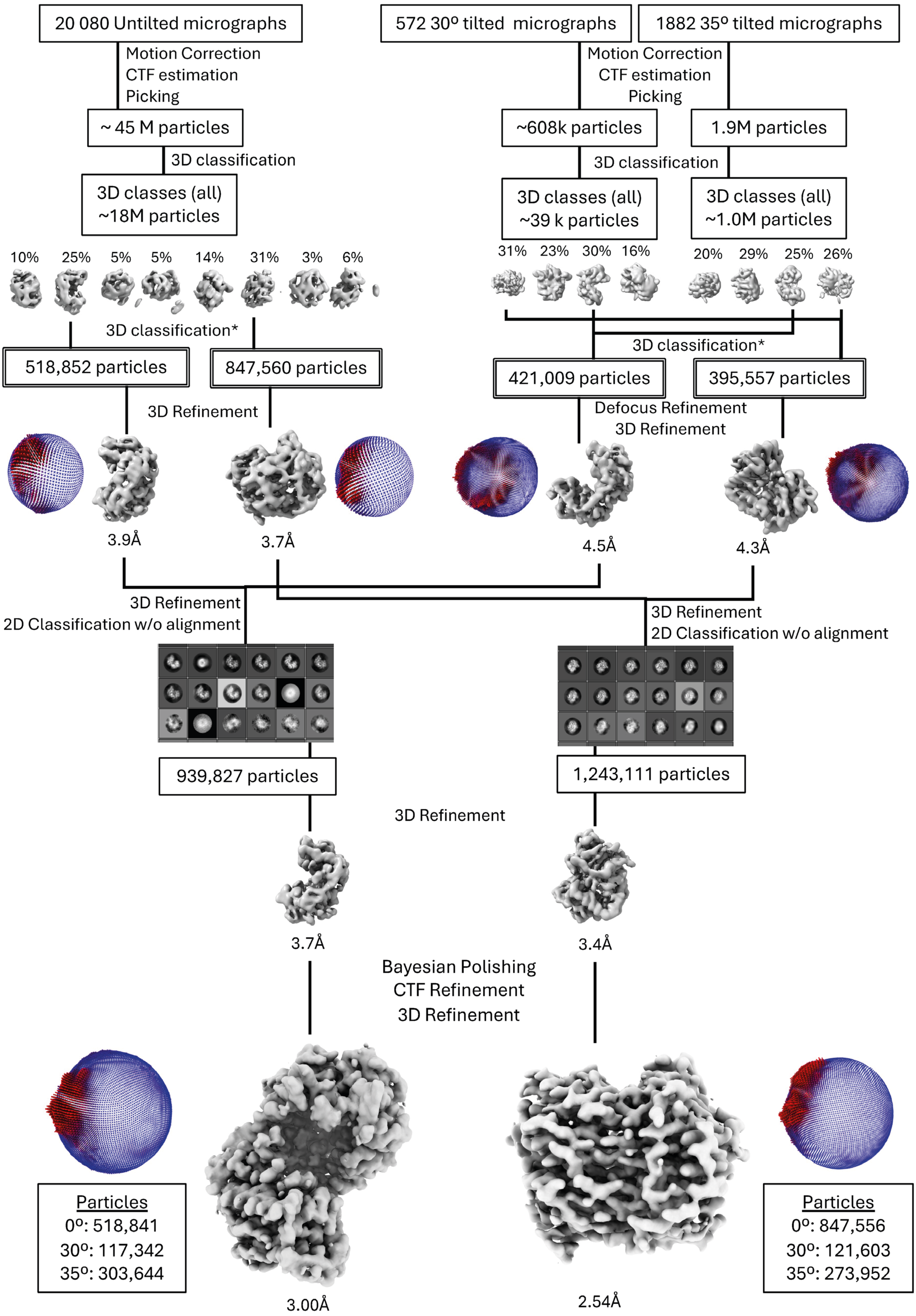
20 080 Untilted micrographs
572 30º tilted micrographs
1882 35º tilted micrographs
Motion Correction
CTF estimation
Picking
Motion Correction
CTF estimation
Picking
~ 45 M particles
~608k particles
1.9M particles
3D classification
3D classification
3D classes (all)
~18M particles
3D classes (all)
~39 k particles
3D classes (all)
~1.0M particles
10% 25% 5% 5% 14% 31% 3% 6%
31% 23% 30% 16%
20% 29% 25% 26%
3D classification*
3D classification*
518,852 particles
847,560 particles
421,009 particles
395,557 particles
3D Refinement
Defocus Refinement
3D Refinement
3.9Å
3.7Å
4.5Å
4.3Å
3D Refinement
2D Classification w/o alignment
3D Refinement
2D Classification w/o alignment
939,827 particles
1,243,111 particles
3D Refinement
3.7Å
3.4Å
Bayesian Polishing
CTF Refinement
3D Refinement
Particles
0º: 518,841
30º: 117,342
35º: 303,644
3.00Å
2.54Å
Particles
0º: 847,556
30º: 121,603
35º: 273,952

Figure S13
